# Mosaic midbrain organoids: a new tool to study Progressive Supranuclear Palsy and advancing clinical neurology research

**DOI:** 10.1101/2024.06.03.597136

**Authors:** Elvira Immacolata Parrotta, Valeria Lucchino, Clara Zannino, Desirèe Valente, Stefania Scalise, Giorgia Lucia Benedetto, Maria Roberta Iazzetta, Mariagrazia Talarico, Francesco Conforti, Silvia Di Agostino, Alessandro Fiorenzano, Aldo Quattrone, Giovanni Cuda, Andrea Quattrone

## Abstract

Progressive supranuclear palsy (PSP) is a severe neurodegenerative disease pathologically characterized by intracellular tangles of hyperphosphorylated tau protein, widely distributed across the neocortex, basal ganglia, and midbrain. Developing effective drugs for PSP presents challenges due to its complex underpinning mechanism and the absence of robust human models that accurately recapitulate biochemical and pathological features of the disease phenotype. Brain organoids have recently emerged as a three-dimensional tissue culture platform to study brain development and pathology. Here, we present a novel induced pluripotent stem cell (iPSC)-derived mosaic midbrain organoid (mMOs) system from four patients with progressive supranuclear palsy-Richardson syndrome (PSP-RS), aimed at reproducing key molecular disease features while reducing variability across organoids derived from different iPSC donors. The PSP-RS 3D model exhibited accumulation of hyperphosphorylated tau protein, predominance of 4R-tau, increased GFAP-positive cells, and PSP-associated histological alterations compared to organoids derived from healthy donors. Pathologically, diseased mMOs showed typical neurofibrillary tangles and tufted-shaped astrocytes, and poorly branched processes of Tyrosine Hydroxylase-immunoreactive cells with thin terminal branches. Our results suggest that mMOs represent a valuable experimental model for PSP research and hold great promise for future identification of new therapeutic targets for progressive supranuclear palsy.

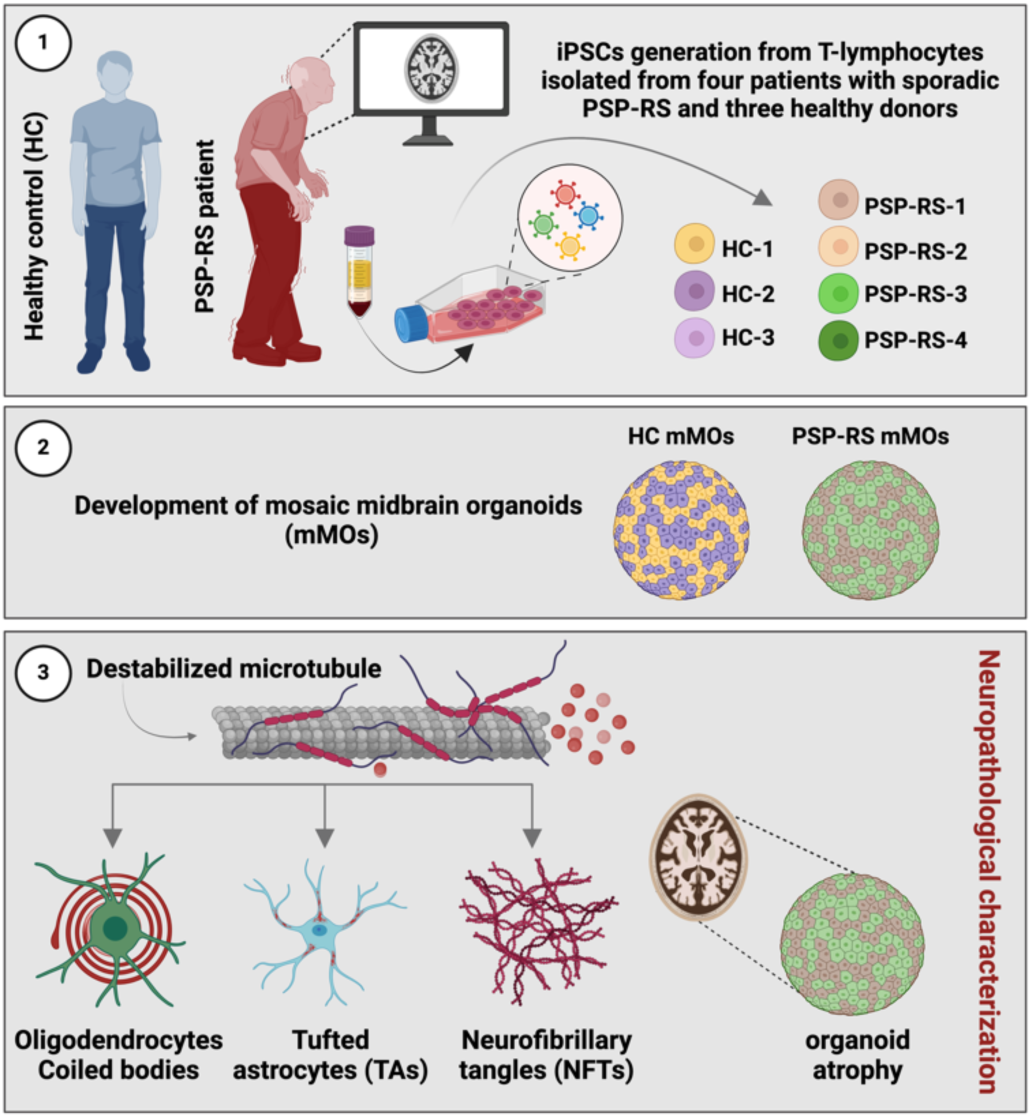

## Introduction

Progressive Supranuclear Palsy (PSP) is a neurodegenerative disorder pathologically characterized by the accumulation of four-repeat (4R) tau isoform in the brain, with the hallmarks of neurofibrillary tangles (NFTs) in neurons, tufted astrocytes (TAs), and oligodendroglial coiled bodies (CBs) ^1–4^. The most common clinical phenotype of PSP, known as Richardson syndrome (PSP-RS), is characterized by vertical oculomotor dysfunction and postural instability as core features. It is often accompanied by akinetic-rigid parkinsonism and cognitive deficits, making it a clinical syndrome highly specific for PSP pathology ^5^. The majority of PSP cases are sporadic, with the most common genetic risk factors being variants in the microtubule-associated protein tau (*MAPT*) ^6–8^. Tau protein is primarily located in the axons of mature neurons where it is involved in stabilizing microtubules and facilitating axonal transport ^9^. The *MAPT* gene undergoes alternative splicing leading to the formation of isoforms containing three microtubule-binding repeats (3R tau) and isoforms containing four microtubule-binding repeats (4R tau) ^10^.

PSP-RS belongs to a group of diseases known as ’4R tauopathies’ due to its predominant shift toward the 4R tau isoform ^11^. In PSP-RS, the basal ganglia and brainstem regions, particularly the midbrain, are the most affected brain structures, especially during the initial phases of the disease. These areas are characterized by significant neuronal loss due to the toxic accumulation of hyperphosphorylated tau protein, which forms insoluble fibrils ^12,13^. Accumulating evidence indicates that tau seeds have the capacity to self-propagate and gradually disseminate across interconnected brain regions over time ^14^.

Despite numerous recent clinical trials testing new disease-modifying drugs targeting tau protein in PSP-RS, none have met primary endpoints ^15^. Nevertheless, they have provided valuable insights into the areas that require attention to improve future research and drug development efforts in PSP. Current biological knowledge of PSP has primarily relied on postmortem brain tissues, thus focusing on the late stage of the disease and lacking systematic investigation into the early-stage PSP pathophysiology. Consequently, a significant gap exists in our understanding of disease progression, particularly in its initial phases, impeding the development of targeted therapies that could potentially intervene in early PSP. Thus, there is an urgent need for model systems that accurately replicate the complex molecular and cellular environment of PSP. The induced pluripotent stem cell (iPSC) technology has revolutionized our capacity to recapitulate inaccessible cell types, offering valuable insights into the mechanisms of various diseases, especially neurological disorders ^16–18^. iPS-derived cerebral organoids have emerged as a powerful tool able to recapitulate architectural and functional features of the human brain, offering a robust *in vitro* model for studying neurological disorders ^19,20^ also at early stages of the disease. Nevertheless, achieving consistent experimental reproducibility is hindered by the differentiation of multiple human iPSC lines, which is complicated by factors such as donor variability, genetic stability, and experimental discrepancies ^21^. Addressing these challenges remains crucial to ensure reliable disease modeling. In this study, we present an innovative approach to mitigate genetic and experimental variations among iPSCs derived from multiple donors. Specifically, we established “mosaic” brain organoids, a highly reproducible, multi-donor human midbrain organoid model developed through the co-development of iPSCs from four individuals affected by PSP-RS within a single organoid. To assess the PSP-RS mosaic model’s efficacy, we conducted a comprehensive side-by-side comparative analysis, examining the biochemical and pathological features of the diseased mMOs against those generated from three age- and gender-matched healthy control individuals.

## Results

### MOSAIC MIDBRAIN ORGANOIDS RECAPITULATE KEY FEATURES OF MIDBRAIN CYTOARCHITECTURE DEVELOPMENT

iPSC cells obtained from four patients diagnosed with sporadic PSP-RS (**Table 1**) were utilized, alongside iPSC cells derived from three healthy control (HC) individuals. **Supplementary** Fig. 1 describes the generation and characterization of iPSC. Data relative to the third iPSC control line are shown in ^22^. Three-dimensional cultures were produced using a pre-established protocol for organoid formation ^23^ with significant modifications made to the initial phase of embryoid body (EBs) formation. Specifically, to minimize variations or discrepancies in organoid differentiation across iPSC batches and donors, we adopted a strategy involving the development of 3D multi- donor mosaic midbrain organoids (mMOs) by pooling iPSCs from four PSP-RS individuals during the embryoid body formation stage. This approach was similarly applied to iPSCs from three healthy control donors (**Fig. 1a**). Prior to initiating the generation of mosaic organoids, we conducted an assessment of each individual iPSC line’s capacity to effectively contribute in midbrain organoid formation (**Supplementary** Fig. 2a). Specifically, we evaluated the differentiation potential of individual organoids by analyzing the expression of early midbrain markers, namely FOXA2 and ZO1 (**Supplementary** Fig. 2b), FOXA2 and LMX1A (**Supplementary** Fig. 2c), NGN2 and ZO1 (**Supplementary** Fig. 2d), along with EN1 (**Supplementary** Fig. 2e), employing immunofluorescence techniques. mMOs were subjected to analysis at various time points over a 120-day period, revealing a consistent and uniform progression of 3D cultures towards the maturation of dopaminergic (DA) neurons. From day -3 to 90 of differentiation, bright field imaging analysis indicated that the midbrain organoids developed into typical morphological structures (**Fig. 1b**). By around day 120, mMO began to exhibit neuromelanin pigmentation, resembling A9 neurons commonly observed within the substantia nigra pars compacta (SNc) region of the ventral midbrain (VM) ^24^. The presence of pigments within mMOs was confirmed through Fontana-Masson staining, known for its specificity in detecting neuromelanin characterized by its distinct dark granular appearance (**Fig. 1b, last image**). The efficiency of midbrain differentiation was assessed by analyzing the expression of early midbrain markers such as *CYNPY1, CORIN, EN1, FOXA2, LMX1A/B, SHH*, and *TH*, as well as the downregulation of the pluripotency-associated genes *OCT4* and *NANOG*, on day 20 (**Fig. 1c**), while late midbrain markers, including *TH, NURR1, DDC, GIRK2, CALB1,* and *DAT,* were assessed on day 60 (**Fig. 1d**). The expression of dopaminergic transcription factors, including FOXA2, LMX1A, and OTX2, along with the tight-junction marker ZO1 (zonula occludens 1) and the microtubule-associated protein 2 (MAP2), involved in microtubule assembly, was validated through immunostaining at an early stage of organoid formation (day 20) (**Fig. 1e, upper panel**). TH-expressing neurons were detected by day 30 (**Fig. 1e, middle panel**), whereas the expression of G-protein-regulated inwardly rectifying potassium channel 2 (GIRK2) and calcium-binding protein 1 (CALB1) was observed by day 60. Additionally, Dopa decarboxylase (DDC), a late midbrain marker, was detected by day 90 of mMO maturation (**Fig. 1e, lower panel**). Expression analysis conducted via quantitative RT-PCR (qPCR) by day 120 unveiled significant upregulation of genes linked to PSP, such as *MAPT*, *STX6, MOBP*, and *PERK* ^7^, in diseased cells compared to healthy controls (**Fig. 1f**). All PSP-RS patients exhibited a homozygosity for the common *MAPT* haplotype H1, which is associated with an increased risk of various tauopathies (**Fig. 1g**). Overall, these data suggest the successful generation of mature dopaminergic neurons within the mMO models, along with the emergence of molecular pathological features.

**Figure 1.**
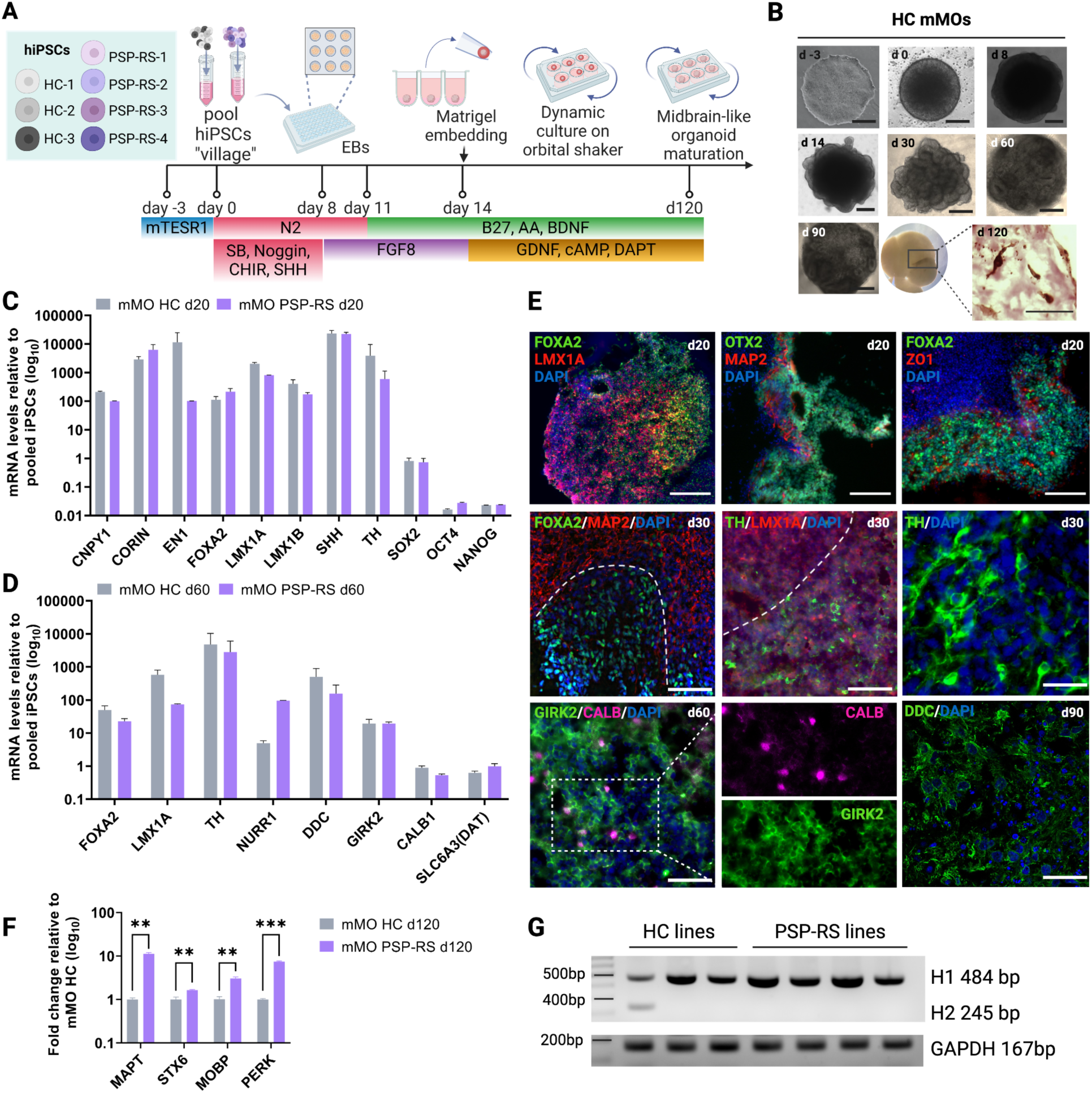
Generation of multi-donor ‘mosaic’ midbrain organoids. a) Graphical representation outlining the progression and timing of mMO differentiation (strategy adapted from Sozzi, E., Nilsson, F., Kajtez, J., Parmar, M., & Fiorenzano, A. (2022). Current Protocols, 2, e555. doi:10.1002/cpz1.555). **b)** Representative bright-field images capturing the progression of hiPSC culture and mMOs differentiation across various time points (from day -3 to day 90) are presented. Scale bar 100 μm. A representative bright-field image of neuromelanin-pigmented mMO from the control group at day 120 is displayed. Scale bar of 100 μm. On the left side, Fontana-Masson staining on sections from HC mMO highlights the release of neuromelanin. Scale bar of 60 μm (bottom right). **c) and d)** RT-qPCR analysis of selected early midbrain and pluripotency markers at differentiation day 20, and late midbrain markers at differentiation day 60, respectively. Values are shown as fold change relative to undifferentiated pooled iPSCs. Statistical analysis was performed using two-tailed unpaired t-test (n=3 mMOs). **e)** Immunostaining for FOXA2/LMX1A, OTX2/MAP2, and FOXA2/ZO1 at day 20 of differentiation (scale bar 200 μm, upper panel); FOXA2/MAP2, TH/LMX1A at day 30 (middle panel); GIRK2/CALB (day 60) and DDC (day 90). Scale bar 50 μm (bottom panel). Nuclei were stained with DAPI. **f)** RT-qPCR analysis of genes associated with PSP neurodegeneration (day 120). Values are given as fold change relative to HC vMOs; statistical analysis was performed using two-tailed unpaired t-test (n=3), ** p ≤ 0.01, *** p ≤ 0.001. Data are presented as mean ± standard error (SEM). **g)** Genotypification of MAPT H1/H2 haplotype using end-point PCR. GAPDH was used as loading control.

**Table 1:** Demographic, clinical and imaging data of patients with progressive supranuclear palsy and controls.

| Data | PSP_001 | PSP_002 | PSP_003 | PSP_004 | HC_001 | HC_002 | HC_003 |
| --- | --- | --- | --- | --- | --- | --- | --- |
| Age at examination | 62 | 62 | 77 | 68 | 54 | 64 | 76 |
| Gender | M | M | F | F | M | M | F |
| Age at disease onset | 58 | 59 | 71 | 62 | / | / | / |
| Disease duration (years) | 4 | 3 | 6 | 6 | / | / | / |
| Diagnosis | Probable PSP-RS | Probable PSP-RS | Probable PSP-RS | Probable PSP-RS | / | / | / |
| PSPRS score | 44 | 25 | 61 | 72 | / | / | / |
| MDS-UPDRS-III | 45 | 53 | 67 | 75 | / | / | / |
| H-Y score | 4 | 4 | 4 | 5 | / | / | / |
| Serum NfL (pg/ml) | 27.4 | 29.5 | 114.0 | 63.3 | 14.2 | 13.5 | 11.1 |
| Midbrain area (mm<sup>2</sup>) | 56 | 71 | 66 | 44 | 129 | 141 | 123 |
| MRPI | 16.47 | 18.08 | 15.20 | 30.08 | 9.94 | 8.02 | 10.38 |

Abbreviations: PSP-RS = Progressive Supranuclear Palsy-Richardson's syndrome, PSPRS = PSP rating scale, M= male; F = Female; MDS-UPDRS-III = MDS-Unified Parkinson's Disease Rating Scale; H-Y = Hoehn and Yahr; MRPI = Magnetic Resonance Parkinsonism Index. Midbrain area was measured on the midsagittal MRI slice. MRPI was calculated as (pons area / midbrain area) x (middle cerebellar peduncle width / superior cerebellar peduncle width). The optimal cut-off to differentiate PSP-RS patients from Parkinson's disease patients and healthy controls was $\geq 13.88$ <sup>25</sup>.

### PSP mMOs EXHIBIT NEURONAL DEGENERATION AND ATROPHY

The sizes of mMOs derived from PSP-RS and healthy control (HC) donors were continuously monitored throughout the culture period, extending up to 120 days. Remarkably, we observed a notable reduction in the volume of organoids derived from PSP-RS individuals compared to those originating from control subjects. While control mMOs continued growing in culture, the patient-derived organoids remained stable after day 60. Over time, the size discrepancy between the two groups became increasingly pronounced. Specifically, by approximately day 120, the diameter of control mMOs measured about 1 cm, whereas PSP-RS mMOs were almost half that size (**Fig. 2, a and b**). This observation is consistent with findings from postmortem studies, which have identified midbrain atrophy as a prominent characteristic of PSP-RS ^26–28^.

**Figure 2.**
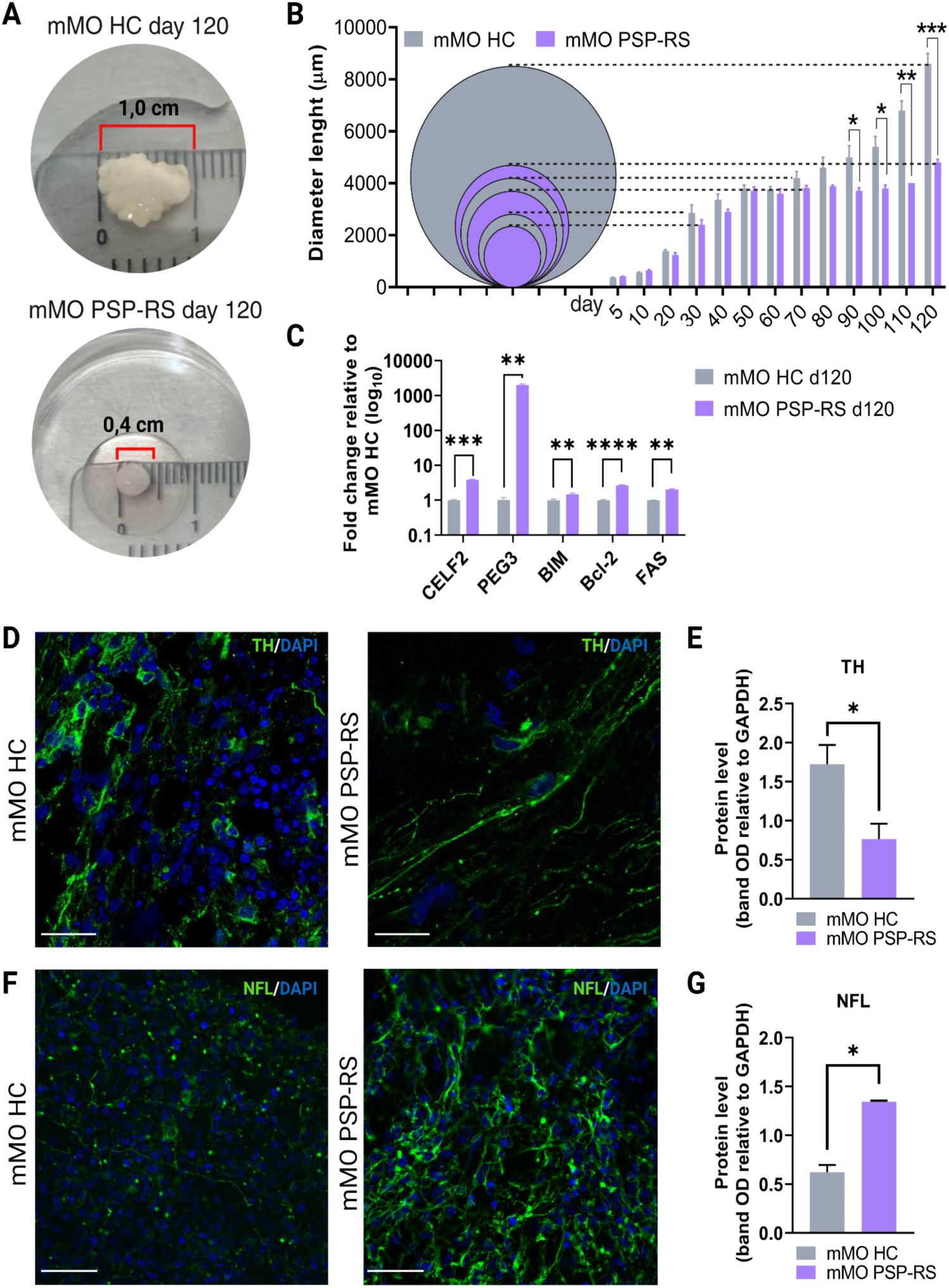
PSP-RS mMOs undergo atrophy and neuronal degeneration. a) PSP-RS mMOs exhibit a remarkable reduction in size, with a diameter of 0.4 cm compared to the 1 cm diameter of HC mMOs. **b)** Growth rates of PSP-RS organoids at various stages are indicated. Error bars represent means ± SEM (n=5). **c)** RT-qPCR analysis of genes associated with apoptosis at day 120 of differentiation. Values are given as fold change relative to HC vMOs; statistic was performed using two-tailed unpaired t-test (n=3), ** p ≤ 0.01, *** p ≤ 0.001, **** p ≤ 0.0001. **d)** Cryosections of HC and PSP-RS mMO at day 120 stained for TH. Scale bar 50 μm. **e)** Quantification of TH protein levels assessed by immunoblot analysis in HC and PSP-RS on day 120 was performed using optical density (OD) measurement. **f)** Cryosections of HC and PSP-RS mMO at day 120 stained for NfL. Scale bar 50 μm. **g)** Immunoblot quantification of NfL protein expression in HC and PSP-RS mMOs at day 120 using OD measurement. Western blots data are shown as mean ± standard error (SEM). Significance was calculated vs. relative HC mMO using t-test, * p ≤ 0.05. GAPDH was used as loading control.

To investigate whether the reduced volume of PSP-RS organoids was associated with cell death under pathological conditions rather than growth arrest, we performed mRNA expression analysis of genes known to trigger apoptosis in neuronal cells, such as *CELF2* and *PEG3* ^29,30^, as well as general apoptotic markers, including *BIM, BCL-2*, and *FAS*. Remarkably, all examined apoptosis-related markers were found significantly upregulated in PSP-RS mMOs compared to HC, providing support for the hypothesis that cell death may contribute to the observed atrophy in PSP-RS organoid (**Fig. 2c**). Furthermore, the processes of tyrosine hydroxylase (TH), a crucial enzyme in the biosynthesis of dopamine and catecholamines in the dopaminergic neurons of the midbrain, were poorly branched and appeared thinner than those of healthy control mMOs (**Fig. 2d**). Furthermore, a notable decrease of TH protein level as quantified through immunoblot analysis was also detected in PSP-RS mMOs (**Fig. 2e and Supplementary** Fig. 3a). These findings are comparable to those occurring in degeneration of dopaminergic (DA) neurons.

Conversely, the expression of neurofilament light chain (NfL), assessed through both immunofluorescence and immunoblot analyses, displayed a significant increase in PSP-RS mMOs compared to HC (**Fig. 2, f and g**, respectively, and **Supplementary** Fig. 3b).

### PSP-RS mMOs EXHIBIT ACCUMULATION OF PHOSPHORYLATED TAU PROTEIN

Pathological aggregates containing hyperphosphorylated tau protein are characteristic features of PSP. To assess whether our 3D model faithfully replicates these aggregations, we quantified protein levels of p-tau Thr231 (**Fig. 3a** and **Supplementary** Fig. 3c), PHF13 recognizing phosphorylated tau at Ser396 (pSer396) (**Fig. 3b** and **Supplementary Fig. d**), p-tau Thr181 (**Fig. 3c** and **Supplementary** Fig. 3e), and AT8 specific for phosphorylated tau at Ser202/Thr205 (**Fig. 3d** and **Supplementary** Fig. 3f) in HC and PSP-RS mMOs at day 60, day 90, and day 120 of differentiation. The analysis revealed a substantial accumulation of these proteins in PSP-RS compared to HC mMOs across all time points, with a peak accumulation observed at day 90, prompting its selection as the optimal time for subsequent analysis. Additionally, immunostaining analysis of PSP-RS and HC mMOs at day 90, labeled with antibodies specific for Thr231, PHF13 (Ser396), Thr181, and AT8, further confirmed a significant accumulation of these proteins in diseased mMOs (**Fig. 3e**). This result was confirmed in mMOs stained at day 120 (**Supplementary** Fig. 3g). PSP is defined as a ‘4R-tauopathy’ due to the predominance of 4R tau isoform present in cytoplasmic inclusions. To investigate the presence of 4R tau, immunostaining was performed using an antibody specific for this isoform. PSP-RS mMO sections exhibited a predominance of 4R tau compared to HC organoids (**Fig. 3f**). Taken together, these results show that our mosaic 3D model of PSP-RS faithfully reproduces the features of tauopathy.

**Figure 3.**
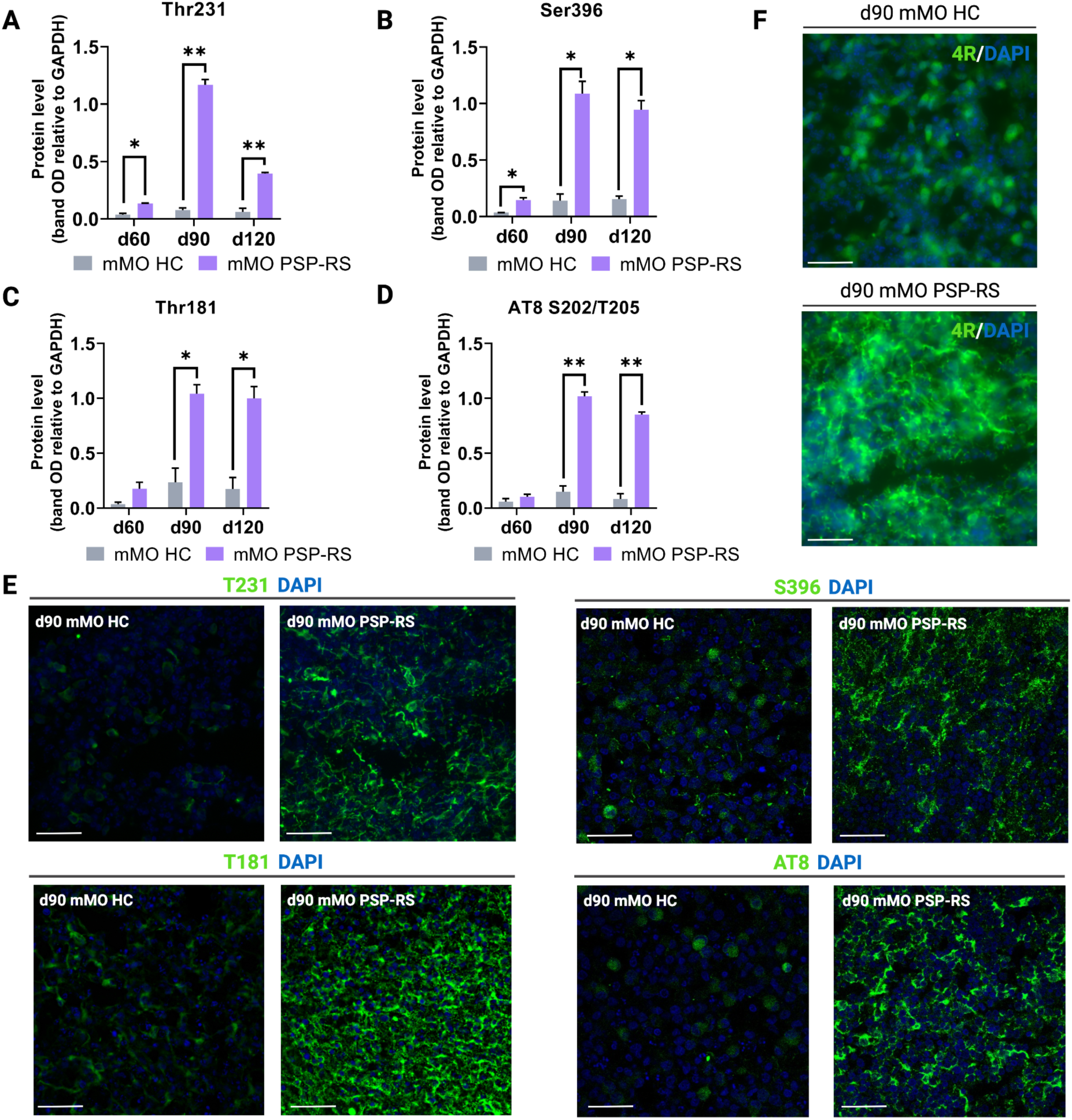
PSP-RS mMOs accumulate hyperphosphorylated tau protein. a-d) Quantification of pThr231 (**a**), pSer396 (**b**), pThr181 (**c**), and AT8 (pSer202/Thr205) (**d)** protein levels at different time points (day 30, day 60, and day 90) assessed using immunoblot analysis in HC and PSP-RS mMOs. Expression levels of pTau were quantified using optical density (OD) measurement. Data are presented as mean ± standard error (SEM). Significance was calculated vs. relative HC mMO using t-test, * p ≤ 0.05, ** p ≤ 0.01. GAPDH was used as loading control. **e)** Representative immunofluorescence images of pThr231, pS396, pThr181, and AT8 labeling in HC and PSP-RS mMOs at day 90. DAPI (blue) was used to stain nuclei. Scale bar 50 μm. **f)** Immunostaining of HC and PSP-RS mMOs cryosection at day 90 with an antibody specific for 4R tau isoform. DAPI (blue) was used for nuclei counterstaining. Scale bar 50 μm.

### HYPERPHOSPHORYLATED TAU ACCUMULATES IN NEURONS AND ASTROCYTES IN ORGANOIDS FROM PSP-RS DONORS

The pathological hallmarks of PSP include the accumulation of four-repeat tau and phosphorylated tau in neurofibrillary tangles (NFTs), frequently exhibiting a globose appearance, and tau-immunoreactive astrocytes (tufted astrocytes, TAs), ^31^ described as part of the degenerative process ^1,2,32^. Here, we investigated the presence of these histological alterations in our 3D model. We first immunoassayed HC and PSP-RS mMO sections with glial fibrillary acid protein (GFAP), and detected a dramatic increase in GFAP-positive cells, reflecting gliosis with immunoreactive astrocytes (**Fig. 4a**). The astroglial population was further confirmed by immunoblot analysis of GFAP protein level indicating a significant and progressive accumulation in diseased organoids assessed at day 120 of maturation (**Fig. 4b**). Subsequently, we performed immunohistochemical (IHC) analysis using Ser396 and AT8 antibodies marking phosphorylated tau protein in NFTs and TA, respectively. We did not detect any positivity for NFTs (S396) or TAs (AT8) in HC mMOs sections (**Fig. 4c and d**). In contrast, PSP-RS mMOs exhibited a strong immunoreactivity for both antibodies reflecting the presence of NFTs and TAs in PSP organoids (**Fig. 4e and f**). Notably, the accumulation of Ser396 tau oligomers was observed in regions where cells exhibited thinner TH branching (**Supplementary** Fig. 4a), suggesting a role of hyperphosphorylated tau in the degeneration of dopaminergic (DA) neurons. Overall, the presence of these histological alterations which are crucial for establishing a pathological diagnosis of ‘definite’ PSP, highlights the strength of our novel 3D model in accurately replicating the key hallmarks of the disease.

**Figure 4.**
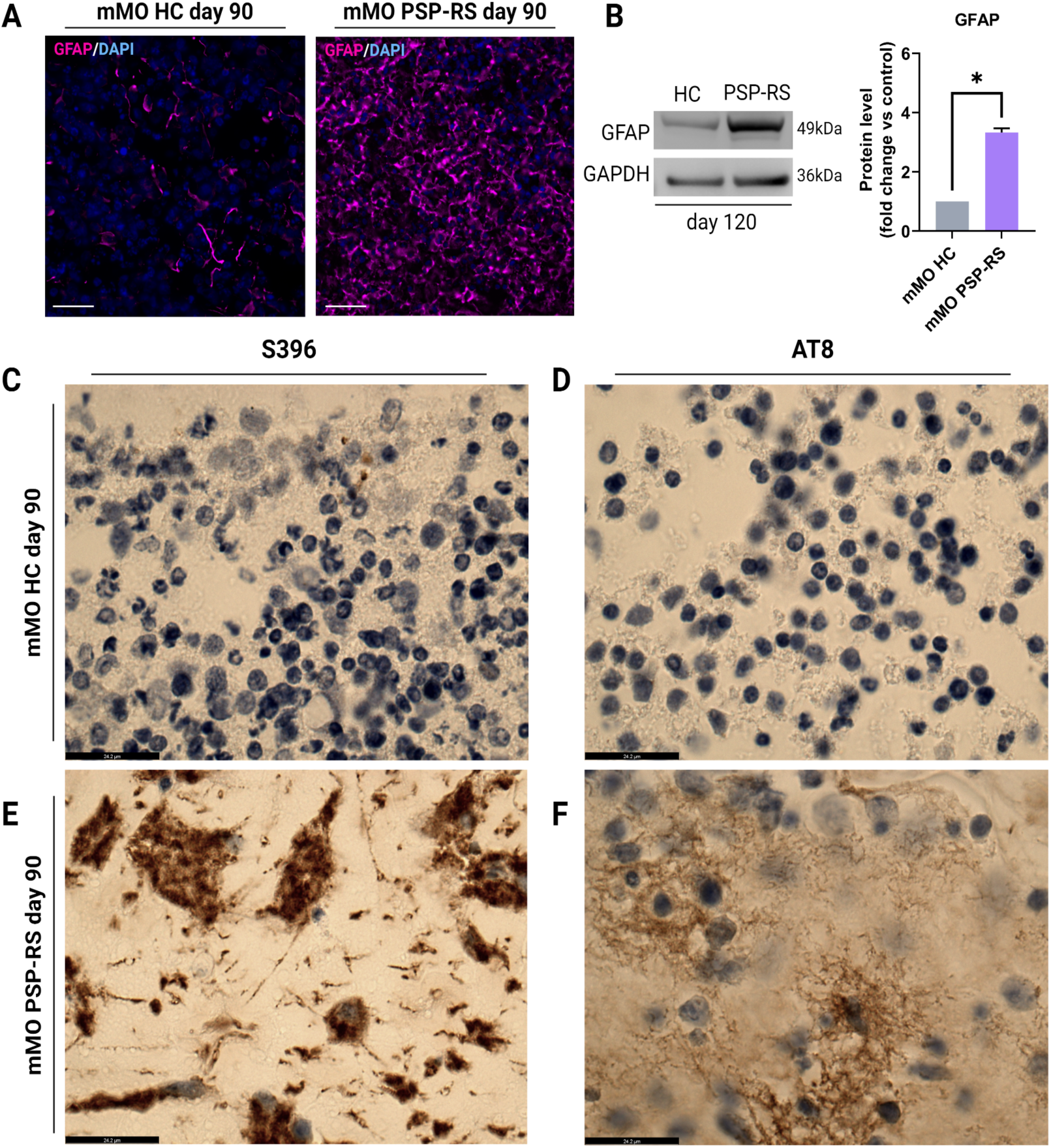
PSP-RS mMOs accumulate GFAP-immunoreactivity and exhibit the presence of neurofibrillary tangles (NFTs) and tufted astrocytes (TAs). a) Immunofluorescence for GFAP in HC and PSP-RS cryosections at day 90. DAPI (blue) was used for nuclear staining. Scale bar 50 µm. **b)** Immunoblot analysis of GFAP protein expression level in PSP-RS and HC mMOs at day 120 and relative quantification conducted using optical density (OD) measurements. Data are presented as mean ± standard error (SEM). Significance was calculated vs. relative HC mMO using t-test, * p ≤ 0.05. GAPDH was used as loading control. **c)** and **d)** Immunocytochemistry for S396 and AT8 (Ser202/Ser205) in mMO sections of HC highlighting the absence of pathological tau aggregates. **e**) and **f)** Immunohistochemistry with DAB counterstained with hematoxylin. Sections were counterstained with hematoxylin. Scale bar 24 µm. **e)** and **f)** Immunocytochemistry for S396 and AT8 in PSP-RS mMOs confirming the presence of neurofibrillary tangles (NFTs) (right) and tufted astrocytes (TAs) (left). Sections were counterstained with hematoxylin. Scale bar 24 µm.

## DISCUSSION

The pathogenesis of neurodegenerative diseases is extremely complex and often associated with impairments in different cellular processes, including oxidative stress ^33^, mitochondrial dysfunction ^34^, response to neuroinflammation ^35^, and toxic protein aggregation ^36^. This complexity makes it challenging to develop effective therapeutic strategies, which are mostly limited to managing symptoms only. Identifying early pathological events and biomarkers responsible for cell degeneration and death would enable the targeting of these critical events to preserve neuronal functionality. The lack of robust *in vitro* models that faithfully replicate the complex cellular processes associated with neurodegeneration has created a bottleneck in the development of novel therapeutics, hindering the discovery of curative treatments ^37–39^. PSP-RS is a rapidly-progressive neurodegenerative disease included among the 4R-tauopathies, with no effective treatment so far, and its exact molecular pathogenesis remains unknown ^40,41^.

Neuropathologically, PSP is characterized by significant neuronal loss, notably affecting the basal ganglia and the midbrain. This loss is attributed to the toxic accumulation of hyperphosphorylated tau protein, which forms insoluble fibrils ^12,13^. Histopathologically, the primary lesions observed in PSP are NFTs and tufted-shaped astrocytes (TAs) ^42^. The predominance of the 4R tau isoform has also been documented ^11^. Three-dimensional human cerebral organoid culture emerges as a novel platform for studying the pathogenesis of neurodegenerative diseases and to evaluate potential therapeutic interventions. In order to better recapitulate the features of PSP-RS and bridge the translational gap between *in vitro* and clinical results, here we present the pioneering *in vitro* midbrain model of PSP-RS. Specifically, we developed mosaic midbrain organoids (mMO) using cells derived from four PSP-RS patients. Our approach offers efficiencies in time and resources compared to individual organoid generation from each donor iPSC line. Importantly, this strategy addresses patient-to-patient variability, ensuring a uniform extracellular environment for all donor iPSCs. We carried out an extensive biochemical characterization allowing us to faithfully reproduce the key neuropathological hallmarks associated with the disease and which have so far been observed in postmortem brain tissues only. First, our findings include a significant reduction in the volume of PSP-RS mMOs, paralleling the atrophy observed in patients’ brains ^27^. The organoid atrophy coincided with the upregulation of genes involved in cell death pathways, suggesting that the observed shrinkage likely results from cell death rather than growth arrest. Our investigation also revealed reduced levels of tyrosine hydroxylase (TH) expression and a poorly branched of TH-positive cells processes appearing thinner in PSP-RS mMOs compared to HC organoids, indicative of DA neuron degeneration in diseased cells. This degeneration was particularly evident in regions with significant accumulation of phosphorylated tau (Ser396). Additionally, we observed a massive presence of gliosis as indicated by the significant increase in the percentage of GFAP-positive in PSP-RS midbrain sections. Finally, we demonstrate the presence of NFTs, oligodendrocytes coiled bodies, and tufted astrocytes (TAs) in PSP-RS. Notably, TAs are currently the most guiding feature, aiding in differentiating PSP from other neurodegenerative diseases. Taken together, our findings indicate that the 3D model herein developed faithfully recapitulates disease-relevant features of PSP, providing a valuable platform for studying the molecular basis underlying its pathogenesis. In addition, midbrain organoids may also serve as a diagnostic tool in the future, allowing to demonstrate the presence of typical PSP histopathological abnormalities *ex vivo*, strongly supporting the clinical diagnosis and helping clinicians in the differential diagnosis of parkinsonism.

### Limitation of the work

One limitation of this work is the lack of cell-type resolved molecular data of PSP-RS. Future investigation in this direction will be focused on an in-depth analysis utilizing single-cell RNA sequencing technology to assess the cell-type composition within the mosaic midbrain organoids and thus elucidating cell-specific gene expression changes and providing a deeper understanding of the intricate network driving PSP progression. However, the development of mMOs has a very high value as a disease model, furthermore the creation of biobanks of tauopathy-specific organoids will facilitate the basic research and the development of precision molecular therapies.

## EXPERIMENTAL DESIGN

### iPSCs generation

After written informed consent, blood samples were collected from four PSP-RS patients and three healthy individuals, who showed no signs of neurological and/or psychiatric illness at the time of sampling. Samples were handled under protocols approved by the University of Catanzaro and the Azienda Ospedaliero-Universitaria ‘Dulbecco’ of Catanzaro. hiPSCs were generated using CytoTuneTM-iPS 2.0 Sendai Reprogramming Kit (Thermo Fisher Scientific) following manufacturer’s instructions with slight modifications. Briefly, PBMCs were cultured in AIM-V medium (Thermo Fisher Scientific) containing 20% FBS, 50 U/mL penicillin, and 50 μg/mL streptomycin (Thermo Fisher Scientific), supplemented with 125 ng/mL Interleukin-2 (IL-2) (R&DSystems). PBMCs were seeded onto CD3-coated dishes (10 μg/mL, BD Biosciences) for T-lymphocytes activation. After 5 days, 5 × 10^5^ T-Lymphocytes were infected with Sendai virus (SeV) at an MOI of 20 in feeder-independent conditions. Three days after transduction, 2x10^5^ cells were plated onto 100-mm dishes coated with Vitronectin (VTN-N) recombinant human protein (Thermo Fisher Scientific, Waltham, MA, USA) in AIM-V medium containing 20% FBS, 50 U/mL penicillin, and 50 μg/mL streptomycin. Approximately, 48 hours after cell seeding, ReproTeSR™ Medium (Stem Cell Technologies, Vancouver, Canada) was progressively added to the cells without medium change. On day 9, medium was completely replaced with ReproTeSR™. Between day 21 and day 28 putative iPS colonies were manually picked and plated on Vitronectin-coated dishes in mTeSR™ Plus (Stem Cell Technologies, Vancouver, Canada) and expanded until tested for SeV loss (around passage 10-12). The absence of chromosomal aberrations in the karyotype of generated iPS lines was confirmed by KaryoStat+ TM Assay (Thermo Fisher Scientific). The characterization of hiPSCs lines used in this study was reported in Supplementary figure S1, except for the HC-1 cell line that was described in a previous work ^22^.

### EBs formation assay

Cells were dissociated into single cells using StemPro Accutase (Thermo Fisher Scientific), then seeded onto low attachment dishes (treated with a solution of 5% poly-2-hydroxyethyl methacrylate in 95% EtOH), and cultured in mTeSR1 medium for seven days. Subsequently, the floating EBs were transferred into plates coated with 5 μg/ml Biolaminin 521LN (Biolamina) and cultured adherently for twenty-one days in DMEM/F12 supplemented with 20% knockout serum replacement (KSR, Thermo Fisher Scientific), 1% Glutamax (Thermo Fisher Scientific), 1% Non-Essential Amino Acids (Thermo Fisher Scientific), 100 μM 2-mercaptoethanol, and 0.5% penicillin and streptomycin.

### Generation of mosaic midbrain organoids

Patient- and healthy donor-derived iPSC lines were cultured in 60-mm dishes pre-coated with Matrigel (Corning, Corning, NY, USA) in mTeSR1 medium (StemCell Technologies, Vancouver, BC, Canada). For the generation of mosaic midbrain organoids (mMOs), iPSCs were detached with Gibco™ StemPro™ Accutase™ Cell Dissociation Reagent (Thermo Fisher Scientific, Waltham, MA, USA) and counted. Cell suspensions derived from the four PSP-RS and three healthy donors were mixed at equal ratios, and 10,000 cells/well were aggregated in ultra-low cell adhesion 96-well plates with U-bottomed conical wells (Corning, NY, USA) to generate mosaic midbrain organoids (mMOs) of PSP-RS and HC, respectively. Pooled cells were cultured mTeSR1 medium supplemented with 10 µM Y-27632 (Miltenyi Biotec, Bergisch-Gladbach, Germany). Midbrain-patterned organoids were generated following the procedure described in ^23^. Briefly, after three days in culture, EBs were transferred to differentiation medium composed of a 1:1 mixture of DMEM/F12 and Neurobasal medium (Thermo Fisher Scientific, Waltham, MA, USA), supplemented with 1:100 N2 supplement, 200 mM L-glutamine, and 10,000 U/mL penicillin/streptomycin (all from Thermo Fisher Scientific, Waltham, MA, USA) supplemented with 10 μM SB431542, 100 ng/ml rhNoggin, 300 ng/ml SHH-C24II, and 1.5 μM CHIR99021 (all from Miltenyi Biotec, Bergisch-Gladbach, Germany). On day 8, the culture medium was supplemented with 100 ng/ml FGF-8b (Miltenyi Biotec, Bergisch-Gladbach, Germany). Starting from day 11, the developing organoids were maintained in Neurobasal medium, enriched with 1:50 B27 supplement (without vitamin A; Thermo Fisher Scientific, Waltham, MA, USA), 200 mM L-glutamine, and 10,000 U/mL penicillin/streptomycin. Additionally, the medium was further supplemented with 100 ng/ml FGF-8b, 20 ng/mL BDNF (Miltenyi Biotec, Bergisch-Gladbach, Germany), and 200 µM L-Ascorbic acid (Sigma-Aldrich, St. Louis, MO, USA). On day 14, organoids were embedded in Matrigel Matrix (Corning, Corning, NY, USA) and cultured in Neurobasal medium supplemented with 1:50 B27 supplement (without vitamin A), 200 mM L-glutamine, and 10,000 U/mL penicillin/streptomycin, supplemented with 20 ng/ml BDNF, 10 ng/ml GDNF (R&D System, Bio-techne, Minneapolis, MN, USA), 200 µM L-Ascorbic acid, 500 µM cAMP (Sigma-Aldrich, St. Louis, MO, USA), and 1 μM DAPT (Tocris, Bio-techne, Minneapolis, MN, USA) to facilitate terminal maturation.

### Immunochemistry (IHC)

For immunohistochemistry, three organoids per group were collected and fixed with 4% paraformaldehyde (Sigma-Aldrich) for 5 hours at room temperature followed by incubation in a 30% w/v sucrose solution (Santa Cruz Biotechnology) overnight at 4°C on an orbital shaker. The next day, the sucrose solution was replaced with a 1:1 mixture of OCT (Avantor) and 30% w/v sucrose and incubated for 6h at 4°C. Organoids were transferred to a cryomold (Avantor) for OCT embedding, and then immediately transferred to dry ice until completely frozen before storage at - 80°C. Sections of 5 µm thickness were made using a cryostat and collected onto Superfrost PlusTM Gold Adhesion microscope slides (Epredia). For staining, the avidin/biotin-based peroxidase Vectastain Elite ABC system and DAB substrate (both from Vector Laboratories) were used. Sections underwent a series of steps to prepare them for IHC. First, they were fixed with a solution of 4% PFA for 10 minutes and incubated with a blocking solution consisting of two drops of Normal serum in PBS^-/-^ for 1 h at room temperature. Subsequently, sections were washed three times with PBS^-/-^ before overnight incubation at 4°C with primary antibody solution. The next day, sections were incubated with a biotinylated secondary antibody for 30 minutes at room temperature, following incubation with the Vectastain solution containing Avidin and Biotinylated Horseradish Peroxidase (HRP) for 30 minutes at room temperature.

Finally, the DAB substrate was added and the reaction was monitored under the microscope. Once the sections turned brown, the reaction was stopped by immersing the slides in distilled water. Nuclei were counterstained with hematoxylin, and the sections were mounted with Eukitt quick-hardening mounting medium. Images were acquired using Leica DM 2500 microscopy. A list of antibodies used for immunostaining is provided in **Supplementary Table 2.**

### Immunofluorescence microscopy

For immunofluorescence analysis, organoids were incubated overnight at 4°C on an orbital shaker in a 30% w/v sucrose solution (Santa Cruz Biotechnology). The next day, the sucrose solution was replaced with a 1:1 mixture of OCT and 30% w/v sucrose and samples were incubated for 6 additional hours at 4°C. Organoids were then transferred to a cryomold (Avantor) and embedded in OCT (Avantor). The cryomold was promptly transferred to dry ice until completely frozen and stored at -80°C. Sections of 20 µm thickness were prepared using a cryostat and collected onto Superfrost PlusTM Gold Adhesion Microscope Slides (Epredia). Sections were fixed with 4% PFA for 10 minutes, then blocked for 1 hour at room temperature with a blocking solution consisting of 5% v/v goat serum (Thermo Fisher Scientific) and 0.3% v/v Triton X-100 (Sigma-Aldrich) in PBS 1X^-/-^. Slides were incubated with primary antibodies overnight at 4°C. The next day, sections were washed three times with PBS for 10 minutes before being subjected to incubation with Alexa Fluor-488, -594, -647 conjugated secondary antibodies (4’,6-diamidino-2-phenylindole, Thermo Fisher Scientific). Afterward, sections were washed again and incubated in a DAPI solution (Thermo Fisher Scientific), followed by mounting using Fluoromount Aqueous Mounting Medium (Sigma-Aldrich). Imaging of the sections was carried out using Leica microscopy systems (Thunder DMi8 and confocal). From each sample, different images were captured. A list of antibodies used for immunostaining is provided in **Supplementary Table 3.**

### Western Blot

For immunoblotting, organoids were lysed by pipetting up and down in RIPA buffer containing 150 mM Sodium Chloride (Sigma-Aldrich), 1% Triton x-100 (Sigma-Aldrich), 0.5% sodium deoxycholate (Sigma-Aldrich), 0.1% Sodium Dodecyl Sulfate (Sigma-Aldrich), 50mM Tris-HCl pH 7.5 (Gibco), supplemented with HaltTM Protease and Phosphatase Inhibitors (Thermo Fisher Scientific). Organoids were then sonicated using a Diagenode Bioruptor (10 cycles, 30 seconds ON/30 seconds OFF) and incubated on ice for 30 minutes. Supernatants were collected by centrifugation at 21,000*g* for 1h at 4°C. The protein content was measured using the Bradford protein assay (Bio-Rad). Equal amounts of proteins (25 µg) were diluted in 1X Bolt^TM^ LDS Sample buffer and BothTM Reducing Agent (Thermo Fisher Scientific). Proteins were denatured for 10 min at 70°C before being resolved in BoltTM 4-12% Bis Tris Plus gels (Thermo Fisher Scientific) using BothTM MES SDS running buffer 20x (Thermo Fisher Scientific) and transferred to nitrocellulose membranes (Bio-Rad) by Trans-Blot TurboTM Transfer System (Bio-Rad). Membranes were blocked for 1 h at room temperature in 5% nonfat dried milk solution (PanReac AppliChem) and incubated overnight at 4°C with primary antibodies. The next day, membranes were incubated with horseradish peroxide (HRP) conjugated secondary antibodies (Jackson ImmunoResearch) at room temperature for 1h. For signal detection, clarityTM ECL (Bio-Rad) was employed and images were captured using AllianceTM Q9-Atom (Uvitec). Densitometry analysis was performed on each band, quantified, and normalized to GAPDH using Image-J software. Subsequently, data were analyzed utilizing one-way ANOVA. The uncropped western blot images are presented in File S1. Antibodies used for immunoblotting are listed in **Supplementary Table 4.**

### DNA extraction and end-point PCR for H1/H2 haplotype

Genomic DNA was obtained from PSP-RS and HC iPS cells using standard phenol-chloroform procedure. Detection of the H1/H2 haplotypes was achieved by determining the 238 bp deletion/insertion variant in intron 9 of the *MAPT* gene using end-point PCR followed by visual 2.5% agarose gel electrophoresis interpretation. Homozygous H1/H1 haplotypes was identified as a single 484 bp band, while the heterozygous H1/H2 haplotype was visualized by the presence of two bands of 484 bp and 246 each. Primers used for end-point PCR are listed in **Supplementary Table 5.**

### RNA extraction, reverse transcription, and quantitative real-time PCR

Total RNA was extracted from each organoid using the TRizol Reagent (Thermo Fisher Scientific), following the manufacturer’s instructions. Subsequently, reverse transcription was performed using the High-Capacity cDNA Reverse Transcription Kit (Thermo Fisher Scientific). Quantitative PCR (qPCR) analyses were conducted in real-time using a QuantStudioTM 7 Pro Real-Time PCR System (Applied Biosystem) and SensiFAST SYBR Hi-ROX Kit (Meridian Bioscience). Gene expression was normalized to the glyceraldehyde 3’-phosphate dehydrogenase (*GAPDH*) housekeeping gene. Analysis was conducted using the comparative Ct (cycle threshold) method. The primer sequences utilized for qRT-PCR analysis are provided in **Supplementary Table 6.**

### Statistical analysis

Statistical analysis was performed using multiple unpaired t tests with Welch correction in GraphPad Prism software, version 9.3.1. Data are represented as the means of two or three biological replicates ± SEM. p values representation: *p < 0.05, **p < 0.01, ***p < 0.001, ****p < 0.0001.

## Author contributions

E.I.P, G.C., A.Q., and A.Q. conceived the study. A.Q. and A.Q. recruited the patients and healthy subjects involved in this study. E.I.P., V.L., C.Z., D.V. performed the experiments and analyzed the data. S.S, G.L.B., M.T. A.F., and M.R.I. contributed to the methodology; S.D.A. and F.C. contributed to immunohistochemical data acquisition and interpretation. E.I.P., G.C., and A.Q. wrote the first draft of the paper and all authors contributed to and approved the final manuscript. G.C. and A.Q. reviewed and edited the final manuscript.

## Materials & Correspondence

Correspondence and request for materials should be addressed to G.C.

## Data and materials availability

All data are available in the main text or the supplementary materials. Any additional information reported in this paper is available from the lead contact upon request.

## Acknowledgments

We thank the patients and healthy subjects in taking part in the study. Without their contributions, this study would not have been possible. We would also like to thank the Centre for Research in Neuroscience and the Institute of Neurology of the University of Catanzaro (Italy) for their collaboration in recruiting patients with progressive supranuclear palsy.

## Funding

PNRR (PIANO NAZIONALE DI RIPRESA E RESILIENZA) National Center for Gene Therapy and Drugs based on RNA technology to GC.

A multiscale integrated approach to the study of the nervous system in health and disease (MNESYS) to GC.

Advanced iPSC-based model of human drug-resistant mesial temporal lobe epilepsy (MTLE) linked to SCN1A mutations (2022J2ARST) - Ministero dell’Università e della Ricerca - PRIN (Progetti di Rilevante Interesse Nazionale) to EIP.

