## Supplementary material for "Mosaic midbrain organoids: a new tool to study Progressive Supranuclear Palsy and advancing clinical neurology research": Supple: Supplementary files_Parrotta et al..pdf

nature neuroscience

**Elvira Immacolata Parrotta *et al.***

**This PDF file includes:**

Figures: S1 to S4

Tables: S1 to S5

**Other Supplementary Materials for this manuscript include the following:**

File S1. Uncropped full length western blots

**Supplementary Fig. 1**

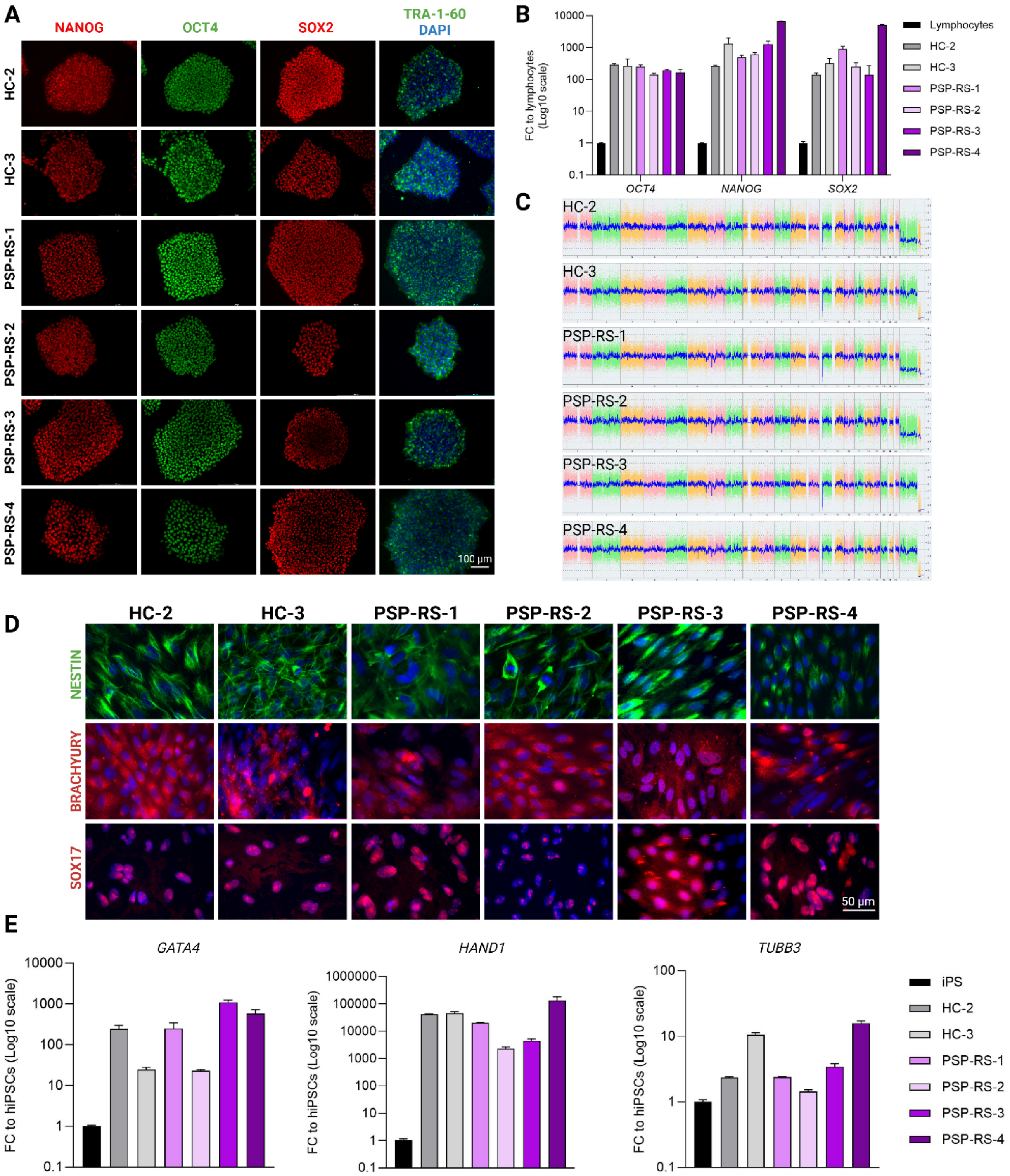

**Supplementary Figure 1. Generation and characterization of iPSC lines. a**) Immunofluorescence analysis revealed a robust expression of pluripotency markers NANOG, OCT4, SOX2, and TRA-1-60 in iPSCs derived from lymphocytes of both PSP-RS patients and healthy control (HC) subjects. Nuclei were counterstained with DAPI (blue). Scale bar 100 µm. **b**) A comparative analysis of mRNA levels demonstrated sustained expression of pluripotency-associated genes *OCT4, NANOG*, and *SOX2* in iPSCs relative to their parental T-lymphocytes cells. **c**) Karyotype analysis using the KaryoStat+™ Assay did not detect any chromosomal aberrations in the generated iPSC lines. **d**) The embryoid body (EB) formation assay displayed positive staining for the lineage-specific markers NESTIN (ectoderm), BRACHYURY (mesoderm), and SOX17 (endoderm), indicating successful differentiation into all three germ layers. **e)** Differentiation markers *GATA4* (endoderm), *HAND1* (mesoderm), and *TUBB3* (ectoderm), were found to be upregulated in EB-derived cells compared to undifferentiated iPSCs, confirming the differentiation potential of iPSCs. Data relative to the third iPSC control line are shown in Parrotta EI, et al., doi: 10.1111/jcmm.14426

**Supplementary Fig. 2**

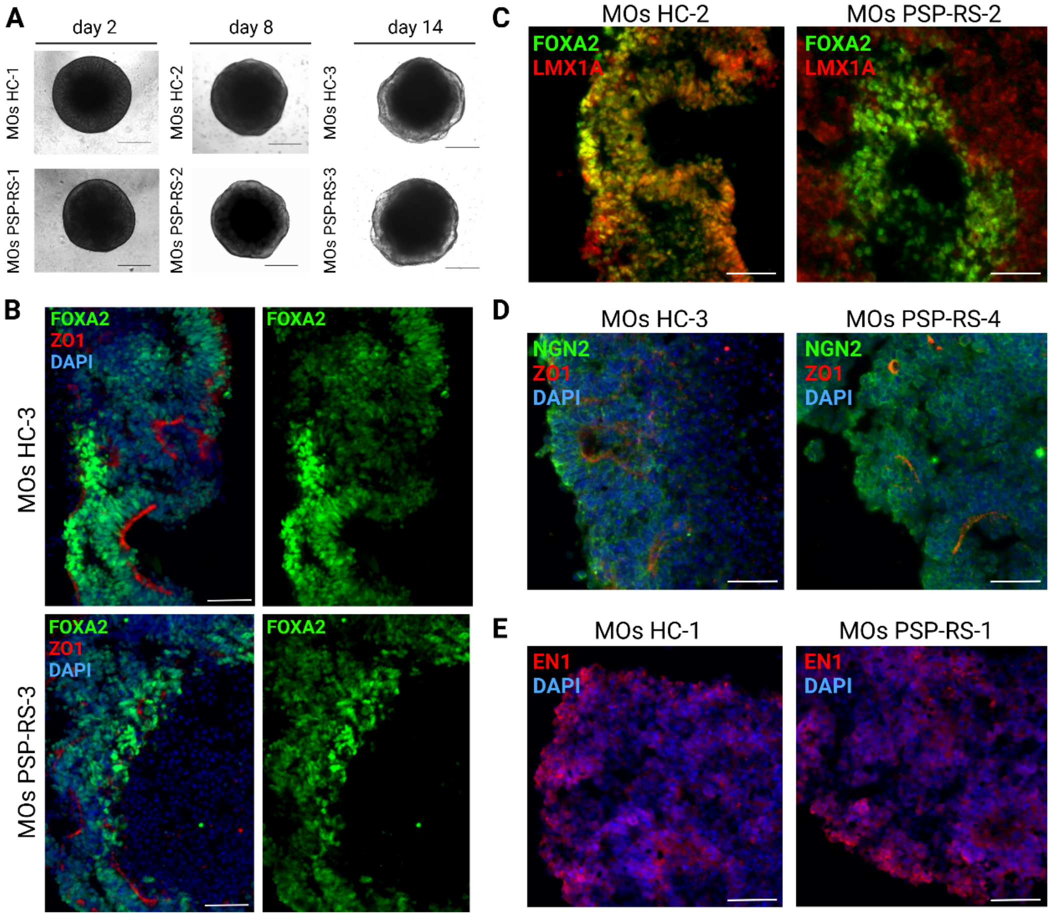


**Supplementary figureF2. Generation of individual midbrain organoids (MOs) from HC and PSP-RS iPS lines a)** Representative bright-field images showing the morphology of MOs at different time points of differentiation. Scale bar: 100 µm. **b-e)** Representative immunochemistry images for early midbrain differentiation markers FOXA2/LMX1A, NGN2/ZO1, FOXA2/ZO1, and EN1 in cryosections of individual HC and PSP-RS MOs at day 15. Nuclei were counterstained with DAPI (blue). Scale bar 50 µm**.**

**Supplementary Fig. 3**

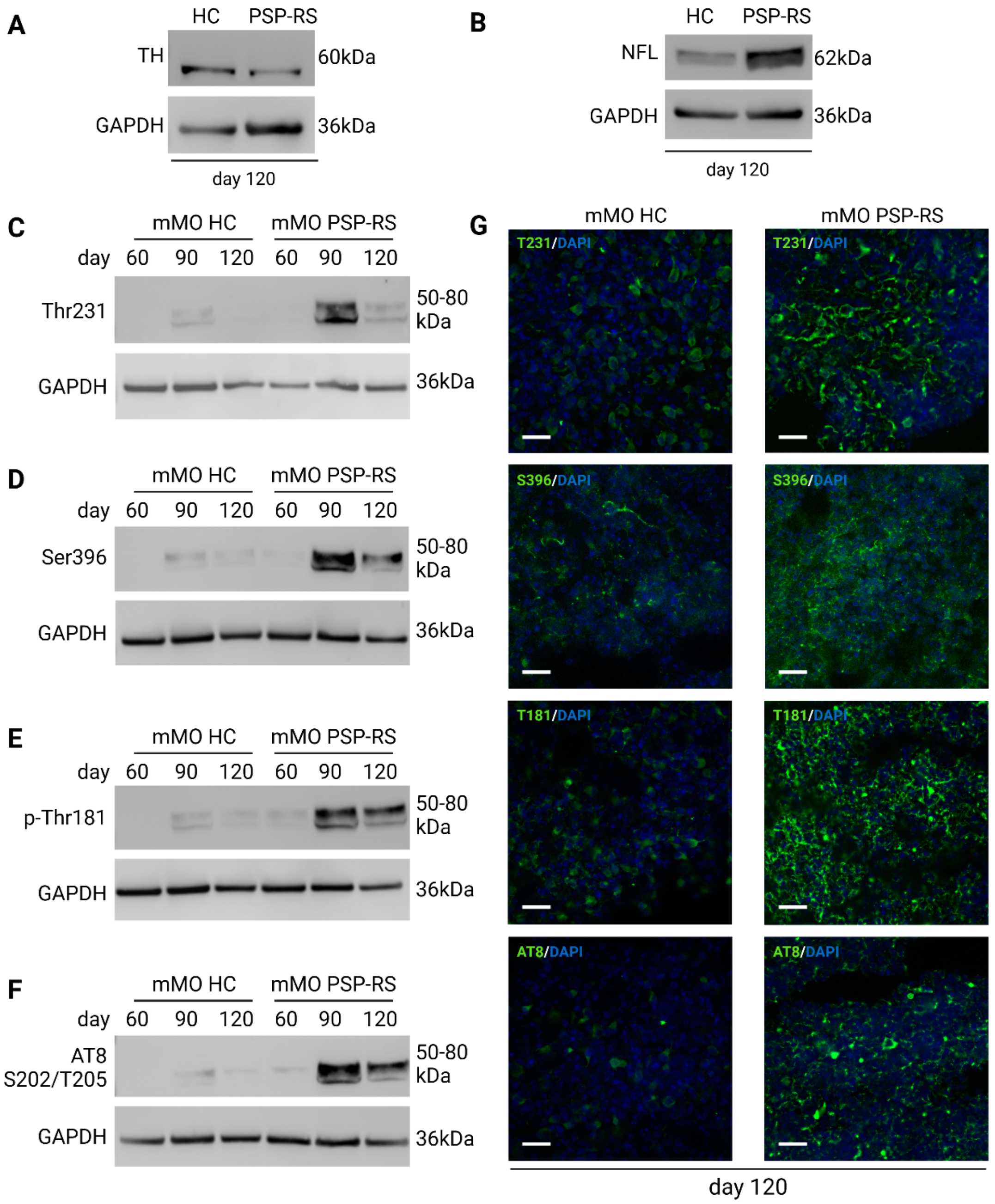


**Supplementary Figure 3. PSP-RS mMOs exhibit increased expression of NfL, decreased expression of tyrosine hydroxylase (TH), and hyperphosphorylated tau accumulation a**) Immunoblot for tyrosine hydroxylase (TH) in HC and PSP-RS mMOs at day 120 of differentiation showing a significant reduction in diseased cells of TH expression. (**b**) Western blot analysis of neurofilament light chain (NfL) in HC and PSP-RS mMOs at day 120 of differentiation highlighting an increased expression of NfL expression in PSP-RS. **c)-f)** Immunoblot analysis to examine the expression levels of phosphorylated tau proteins: pThr231(**c**), pS396 (**d**), pThr181(**e**), and AT8 (**f**), in both HC and PSP-RS mMOs at days 60, 90, and 120. PSP-RS organoids exhibit a progressive accumulation of phosphorylated tau with a peak expression at day 90. GAPDH was used as loading control in all Western Blot analysis. **g**) Immunostaining with phosphorylated tau T231 (top panel), S396 (second panel), T181 (third panel), and AT8 (bottom panel) at day 120 in HC and PSP-RS mMOs. DAPI was used to stain nuclei. Scale bar: 50 µm.

**Supplementary Fig. 4**

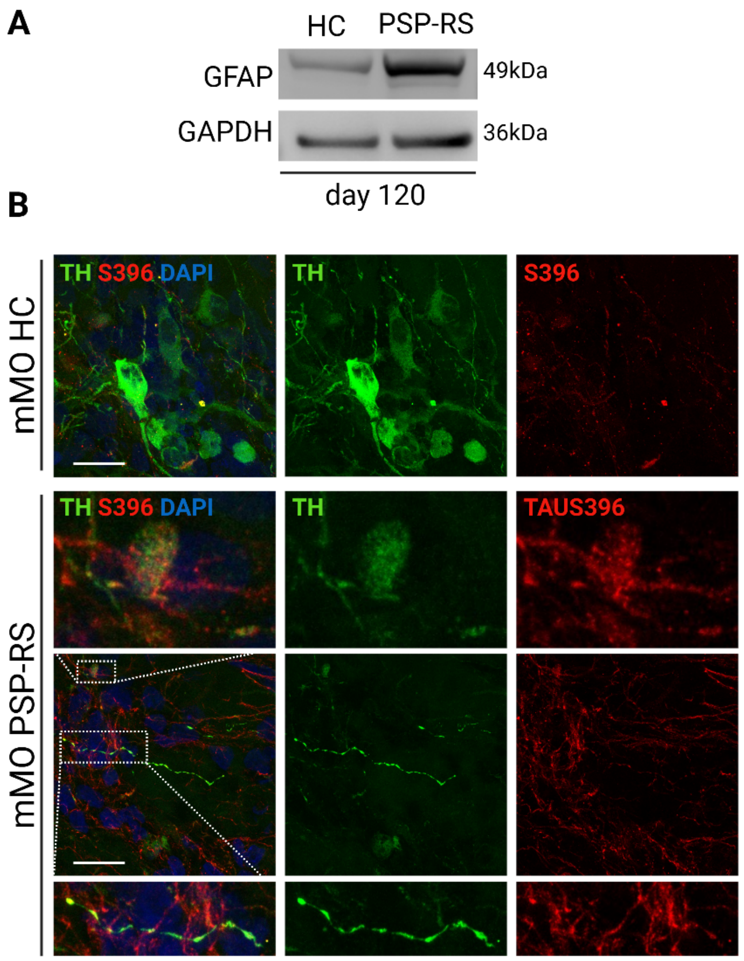


**Supplementary Figure 4. PSP-RS mMOs exhibit an accumulation of GFAP-positive cells and an increase of phosphorylated tau S396 in conjunction with poorly branched TH-immunoreactive cells. a**) Representative immunostaining images at day 120 in HC and PSP-RS mMOs stained with antibodies against Tyrosine Hydroxylase (TH) and S396 revealing in PSP-RS an accumulation of S396 in areas where TH-expressing cells show a thinner branching. DAPI (blue) was used for nuclei stains. Scale bar 50 µm.

**Supplementary Table 1.** List of antibodies used for Immunohistochemistry analysis

| Antibody | Host species | Dilution | Cat. No | Company |
|---|---|---|---|---|
| pTAU (Ser202/Thr205) | Mouse | 1:500 | MN1020 | Invitrogen |
| pTAU (Ser396) | Rabbit | 1:500 | 44752G | Invitrogen |

**Supplementary Table 2.** List of antibodies used for Immunofluorescence analysis

| Antibody | Host species | Dilution | Cat. No | Company |
|---|---|---|---|---|
| FOXA2 | Mouse | 1:1000 | sc-101060 | Santa Cruz |
| OCT4 | Mouse | 1:200 | 75463 | Cell Signaling |
| NANOG | Rabbit | 1:200 | PA1-097 | Invitrogen |
| SOX2 | Rabbit | 1:200 | Ab97959 | Abcam |
| TRA1-60 | Mouse | 1:100 | 41-1000 | Invitrogen |
| NESTIN | Mouse | 1:500 | 60091 | StemCell Tech. |
| SOX17 | Goat | 1:20 | AF1924 | R&D System |
| BRACHYURY | Goat | 1:20 | AF2085 | R&D System |
| FOXA2 | Rabbit | 1:300 | Ab108422 | Abcam |
| LMX1A | Rabbit | 1:200 | PA5-115517 | Invitrogen |
| OTX2 | Goat | 1:2000 | AF1979 | R&D System |
| MAP2 | Chicken | 1:1000 | PA-10005 | Invitrogen |
| ZO1 | Mouse | 1:300 | 33-9100 | Invitrogen |
| TH | Mouse | 1:50 | MA1-24654 | Invitrogen |
| NFL | Rabbit | 1:100 | 2837S | Cell Signaling |
| βTUB III | Mouse | 1:250 | 480011 | Invitrogen |
| CALB | Rabbit | 1:200 | BK13176S | Cell Signaling |
| GIRK2 | Goat | 1:100 | Ab65096 | Abcam |
| DDC | Goat | 1μg/mL | PA547512 | Invitrogen |
| pTAU (Ser202/Thr205) | Mouse | 1:500 | MN1020 | Invitrogen |
| pTAU (Ser396) | Rabbit | 1:500 | 44752G | Invitrogen |
| pTAU (Thr181) | Rabbit | 1:100 | 12885 | Cell Signaling |
| TAU | Mouse | 1:100 | AHB0042 | Invitrogen |
| GFAP | Chicken | 1:1000 | PA1-10004 | Invitrogen |
| pTAU (Thr231) | Mouse | 1:500 | MN1040 | Invitrogen |
| TAU (3R) | Mouse | 1:250 | S-05-803 | Merck |
| TAU (4R) | Rabbit | 1:100 | 79327S | Cell Signaling |

| | | | | |
|---|---|---|---|---|
| TH | Rabbit | 1:100 | BK58844S | Cell Signaling |
| Anti-Mouse IgG Alexa Fluor 488 | Goat | 1:2000 | A-11001 | Invitrogen |
| Anti-Rabbit IgG Alexa Fluor 594 | Goat | 1:500 | A-11012 | Invitrogen |
| Anti-Chicken IgY (H+L)<br>Alexa Fluor 594 | Goat | 1:200 | A-32759 | Invitrogen |
| Anti-Rabbit IgG Alexa Fluor 488 | Goat | 1:500 | A-11008 | Invitrogen |
| Anti-Mouse IgG Alexa Fluor 594 | Goat | 1:200 | A-11005 | Invitrogen |
| Anti-Goat IgG Alexa Fluor 488 | Chicken | 1:500 | A-21467 | Invitrogen |
| Anti-Rabbit IgG Alexa Fluor 647 | Goat | 1:500 | A-21244 | Invitrogen |
| Anti-Mouse IgG Alexa Fluor 647 | Goat | 1:500 | A-21235 | Invitrogen |

**Supplementary Table 3.** List of antibodies used for Western blot analysis

| Antibody | Host species | Dilution | Cat. No | Company |
|---|---|---|---|---|
| TH | Rabbit | 1:1000 | BK58844S | Cell Signaling |
| GFAP | Chicken | 1:5000 | PA1-10004 | Invitrogen |
| NFL | Rabbit | 1:1000 | 2837S | Cell Signaling |
| GAPDH | Rabbit | 1:1000 | bs10900R | Bioss Antibodies |
| pTAU (Thr231) | Mouse | 1:500 | MN1040 | Invitrogen |
| pTAU (Ser396) | Rabbit | 1:1000 | 44752G | Invitrogen |
| pTAU (Thr181) | Rabbit | 1:1000 | 12885 | Cell Signaling |
| pTAU (Ser202/Thr205) | Mouse | 1:500 | MN1020 | Invitrogen |
| Peroxidase AffiniPure Donkey Anti-Rabbit IgG (H+L) | | 1:10000 | 711-035-152 | Jackson Immuno Research |
| Peroxidase AffiniPure Sheep Anti-Mouse IgG (H+L) | | 1:10000 | 515-035-062 | Jackson Immuno Research |

**Supplementary Table 4.** Primers used for MAPT haplotype

| Gene | Primer sequence |
|---|---|
| MAPT haplotype | For GGAAGACGTTCTCACTGATCTG<br>Rev AGGAGTCTGGCTTCAGTCTCTC |

**Supplementary Table 5.** Primers used for quantitative Polymerase Chain reaction (qPCR) analysis

| Gene | Primer sequence |
|---|---|
| CNPY1 | **For** GGAAGACCCTGTGACGAAGG; **Rev** TCCTGGGCGATAAGTGAGGA |
| CORIN | **For** CCCCGGGAAACTGCAATGTA; **Rev** GCGATGCTCTGTTGTGGGAT |
| EN1 | **For** GCCCGTGGTCAAAACTGACT; **Rev** GGAACTCCGCCTTGAGTCTC |
| FOXA2 | **For** CTGGTCGTTTGTTGTGGCTG; **Rev** CGTGTTCATGCCGTTCATCC |
| LMX1A | **For** GCCATCGAGCAGAGTGTCTA; **Rev** TACTGAGGGAGGTGTCGTCG |
| LMX1B | **For** CGGACTGCGCCAAGATGTT; **Rev** TTGACTCGCATCAGGAAGCG |
| SHH | **For** TGGACATCACCACGTCTGAC; **Rev** GGAAGCAGCCTCCCGATTT |
| TH | **For** TGTACTGGTTCACGGTGGAGT; **Rev** TCTCAGGCTCCTCAGACAGG |
| SOX2 | **For** GGGAAATGGGAGGGGTGCAAAAGAGG<br>**Rev** TTGCGTGAGTGTGGATGGGATTGGTG |
| OCT4 | **For** GACAGGGGGAGGGGAGGAGCTAGG **Rev**CTTCCCTCCAACCAGTTGCCCCAAAC |
| NANOG | **For** TGCAAGAACTCTCCAACATCCT **Rev** ATTGCTATTCTTCGGCCAGTT |
| GATA4 | **For** GGCCTGTCATCTCACTACGG; **Rev** ATGGCCAGACATCGCACT |
| HAND1 | **For** CCAGCTACATCGCCTACCTG; **Rev** CCGGTGCGTCCTTTAATCCT |
| βTUB III | **For** GCAACTACGTGGGCGACT **Rev** CGAGGCACGTACTTGTGAGA |
| DDC | **For** GCCGCTATCATGGAGAAGCT; **Rev** AGAAAGGAATCAGGCCAGCC |
| GIRK2 | **For** CACATCAGCCGAGATCGGAC; **Rev** GGTAGCGATAGGTCTCCCTCA |
| CALB1 | **For** ATCCCTCATCACAGCCTCAC; **Rev** TTTGCCCATACTGATCCACA |
| DAT | **For** GTCTGTTTGGATTGACGCGG; **Rev** AAGGAGAAGACGACGAAGCC |
| NURR1 | **For** TGCCGATTTCAGAAGTGCCT; **Rev** CGAGGGCACTGATCAGACTC |
| MAPT | **For** AAGTCGCCGTCTTCCGCCAAG **Rev** GTCCAGGGACCCAATCTTCGA |
| STX6 | **For** TACTCGGCAAGTTGTCAGGG; **Rev** CGGTCCAGACGCCCATATTT |
| MOBP | **For** CCAATCAGCAGATGTGTACGG; **Rev** AGCCGCTCTTACAGATGCTG |
| PERK | **For** AGCCAATTCAATGCCTGGGA; **Rev** TAACAATGCCCGGGTGTTCA |
| CELF2 | **For** GCCAGATAGAAGAATGCCGGA; **Rev** GTGCCATTGCCCTTGTAGA |
| PEG3 | **For** GGGCCACTCATCAAGATCCA; **Rev** TTCCCGATTTGGAACTGCGT |
| BIM | **For** GCTGTCTCGATCCTCCAGTG; **Rev** TCCAATACGCCGCAACTCTT |

| | |
|---|---|
| Bcl2 | **For** TGTGGATGACTGAGTACCTG; **Rev** GCCAGGAGAAATCAAACAGAG |
| FAS | **For** GGAGTACACAGACAAAGCCCA; **Rev** TTTGGTGCAAGGGTCACAGT |

**Supplementary File 1. Uncropped Western Blot images**

The bands shown are results of the immunoblot analysis conducted in this study. The red rectangles highlight the specific bands referenced in the text. For all uncropped images provided, the correspondence to the main or supplementary figure is indicated.

**2) Figure 2e**

1° replicate

d120

HC PSP-RS

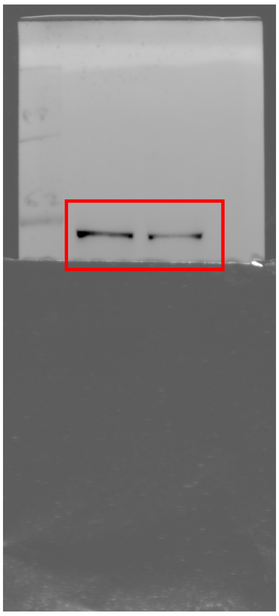

TH MW 55-70 kDa

d120

HC PSP-RS

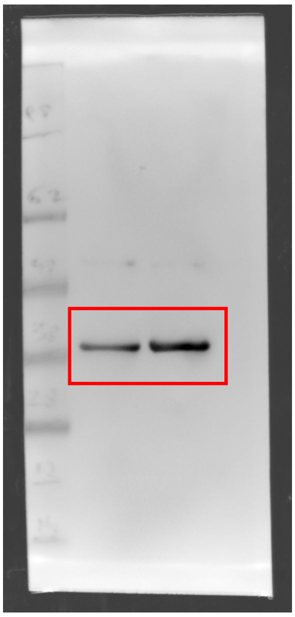

GAPDH MW 36 kDa

2° replicate

d120

HC PSP-RS

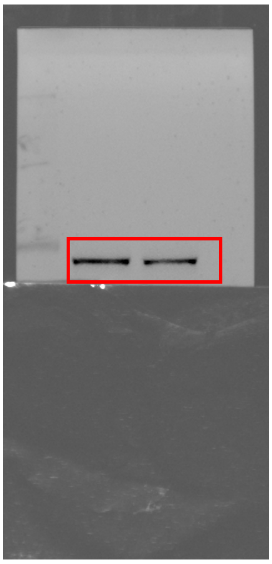

TH MW 55-70 kDa

d120

HC PSP-RS

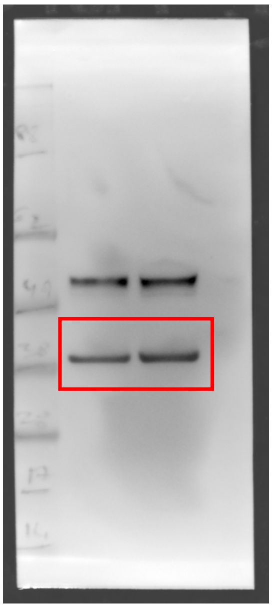

GAPDH MW 36 kD

#### 3) Figure 2g

1° replicate

d120

HC PSP-RS

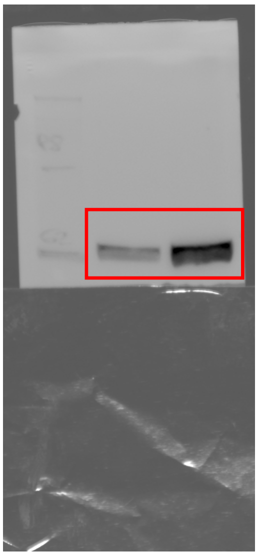

NfL MW 70 kDa

d120

HC PSP-RS

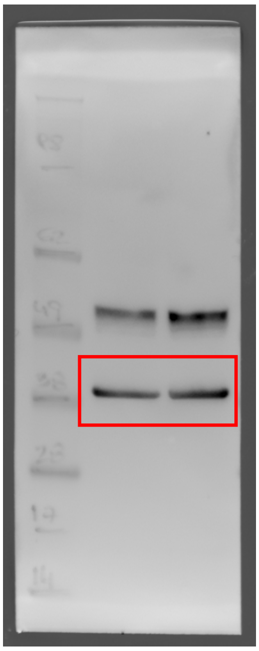

GAPDH MW 36 kDa

2° replicate

d120

HC PSP-RS

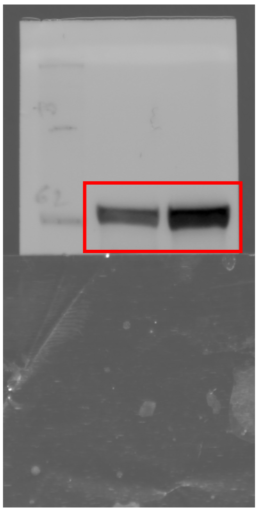

NfL MW 70 kDa

d120

HC PSP-RS

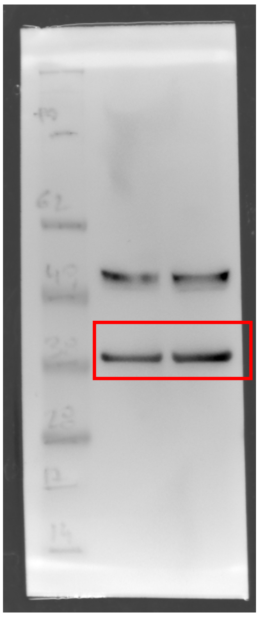

GAPDH MW 36 kDa

### 4) Figure 3a

1° replicate

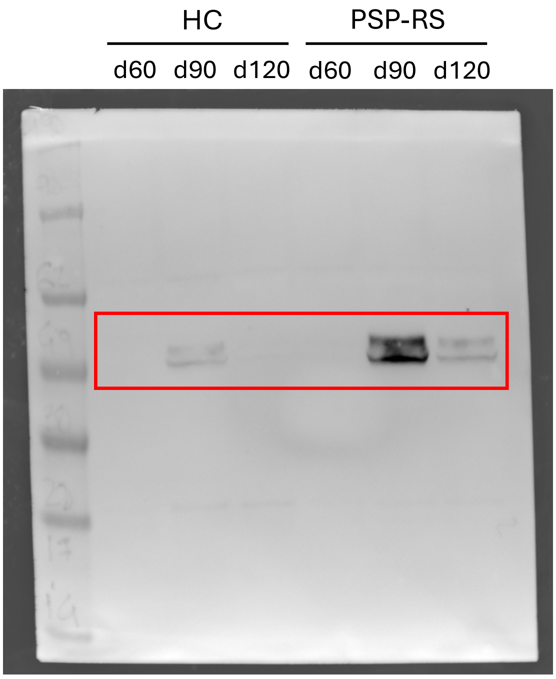


pTAU (Thr231) MW 50-70 kDa

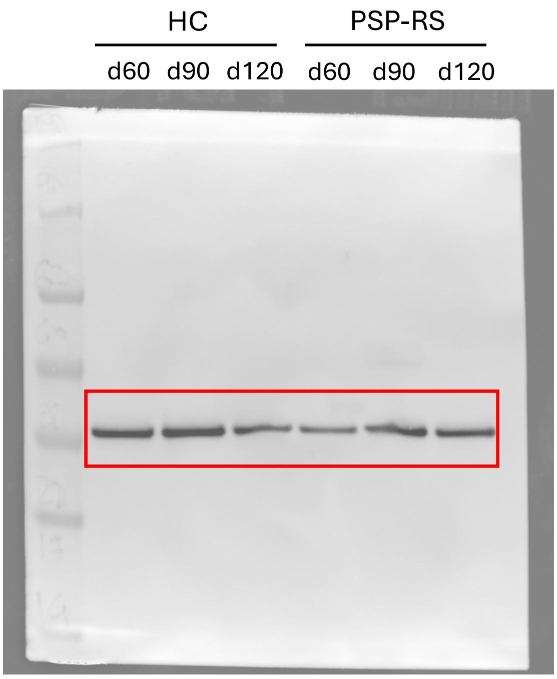


GAPDH MW 36 kDa

2° replicate

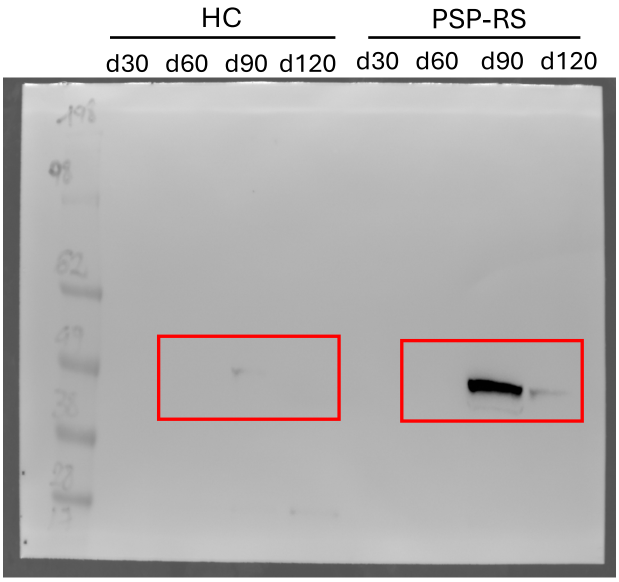


pTAU (Thr231) MW 50-70 kDa

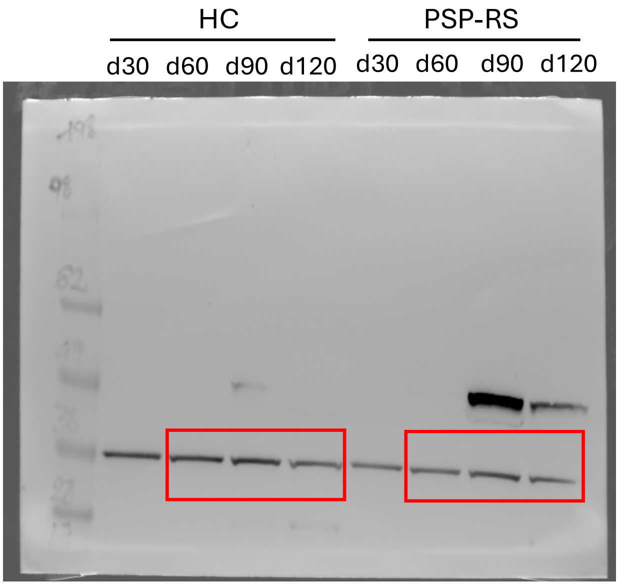


GAPDH MW 36 kDa

### 5) Figure 3b

1° replicate

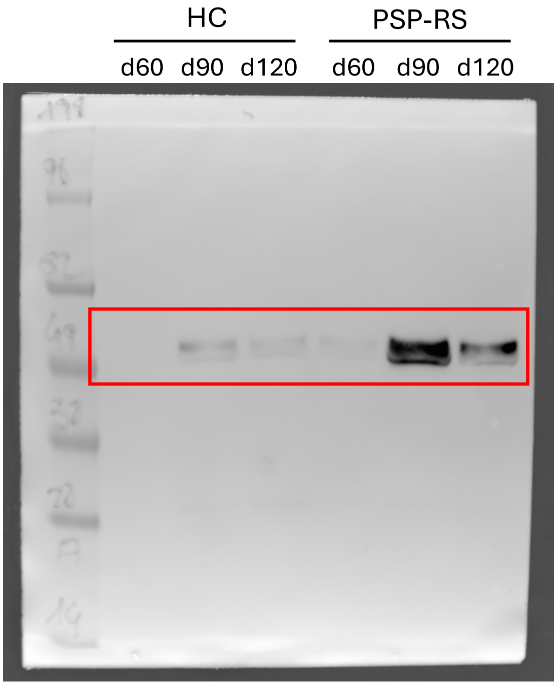


pTAU (Ser396) MW 50-70 kDa

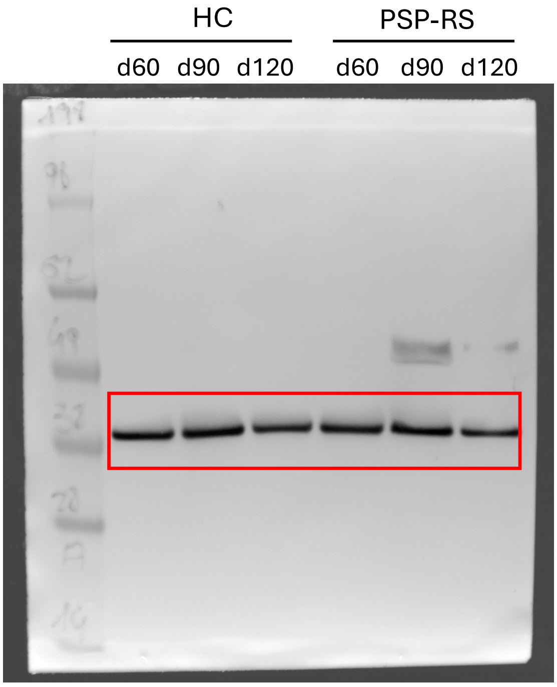


GAPDH MW 36 kDa

2° replicate

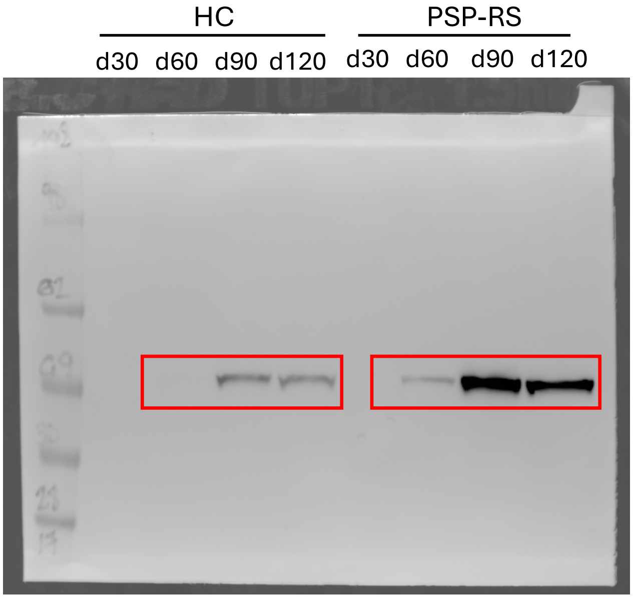


pTAU (Ser396) MW 50-70 kDa

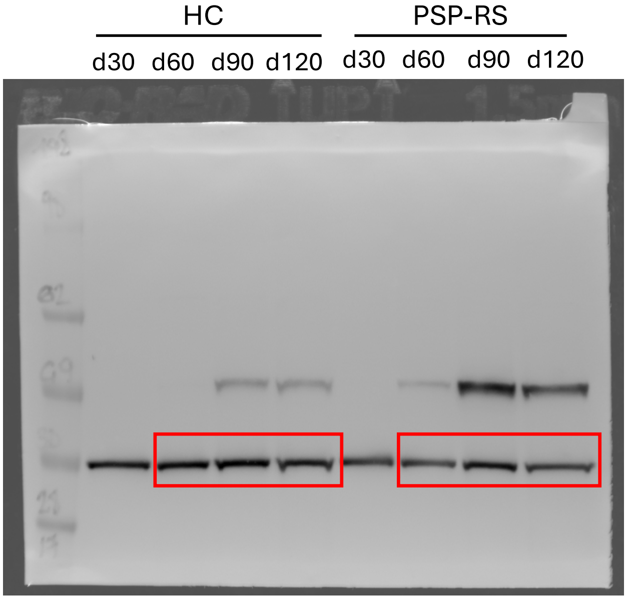


GAPDH MW 36 kDa

### 6) Figure 3c

1° replicate

HC | PSP-RS

d60 d90 d120 d60 d90 d120

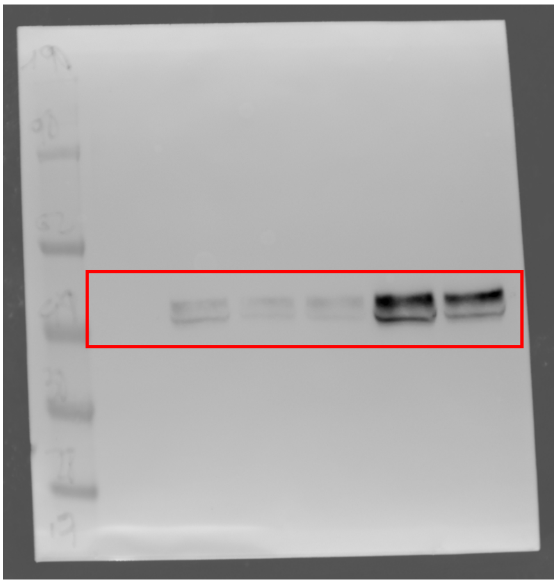

pTAU (Thr181) MW 50-70 kDa

HC | PSP-RS

d60 d90 d120 d60 d90 d120

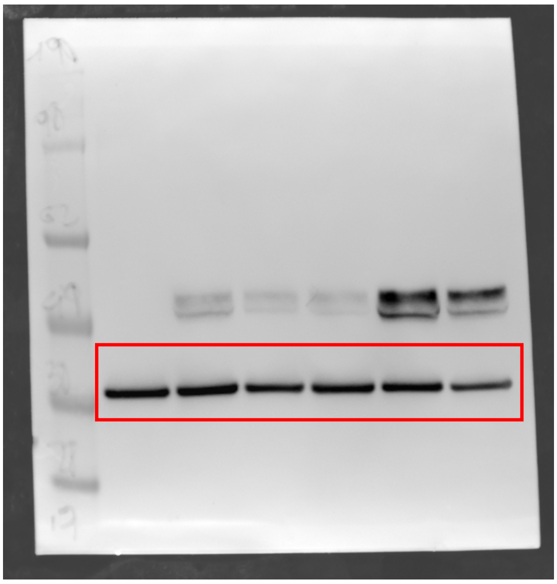

GAPDH MW 36 kDa

2° replicate

HC | PSP-RS

d30 d60 d90 d120 d30 d60 d90 d120

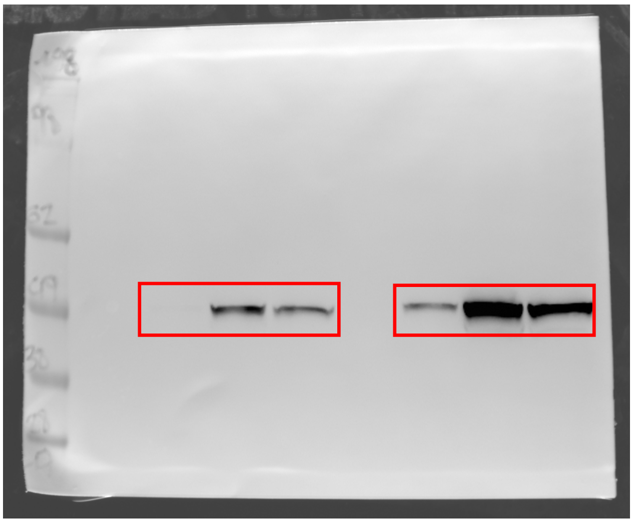

pTAU (Thr181) MW 50-70 kDa

HC | PSP-RS

d30 d60 d90 d120 d30 d60 d90 d120

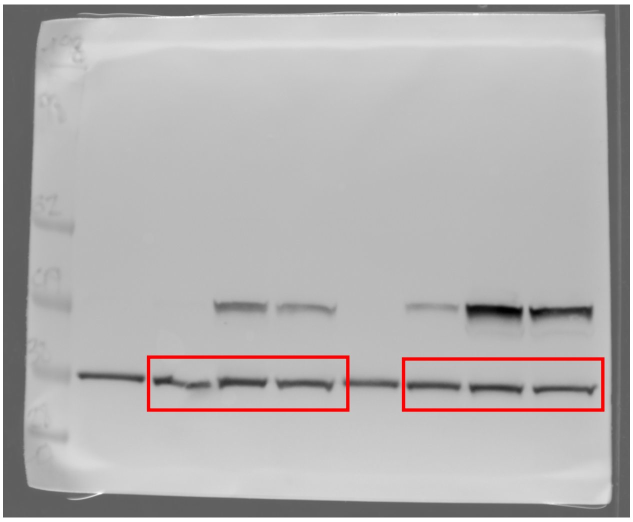

GAPDH MW 36 kDa

#### 7) Figure 3d

1° replicate

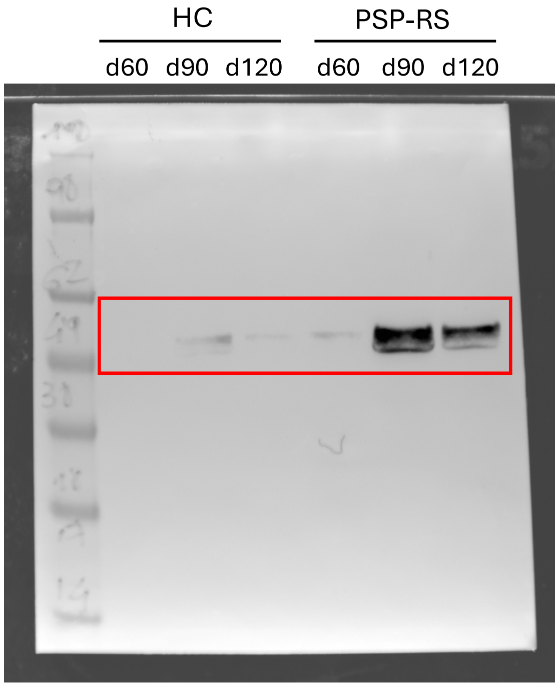


pTAU(Ser202/Thr205) MW 50-70 kDa

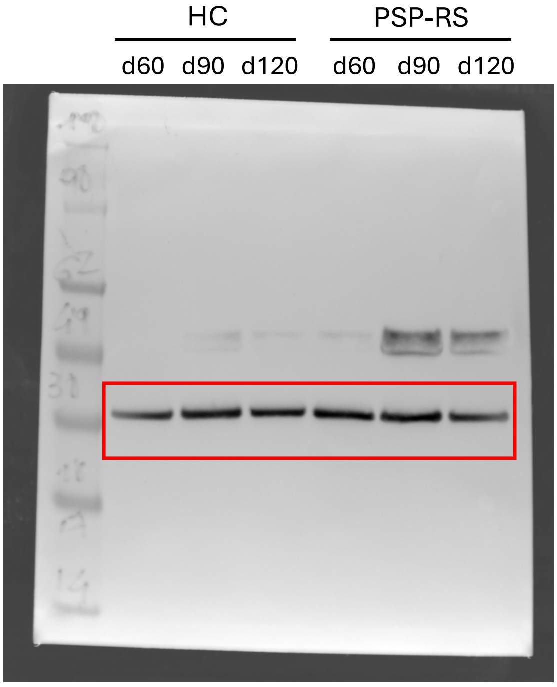


GAPDH MW 36 kDa

2° replicate

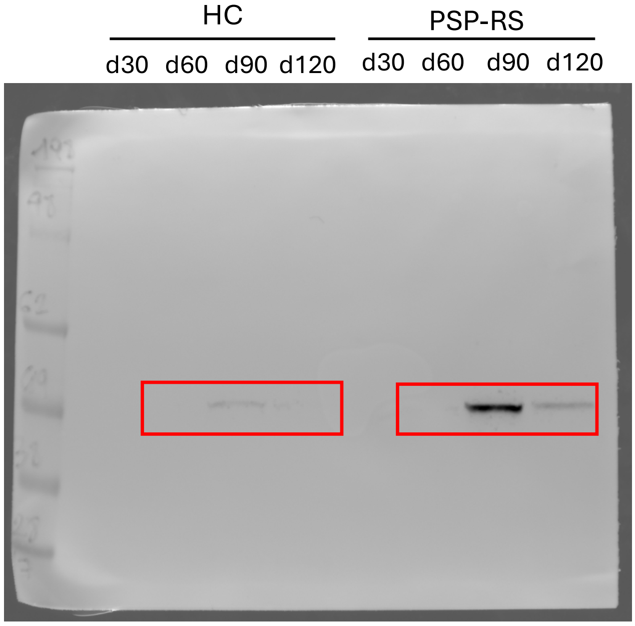


pTAU(Ser202/Thr205) MW 50-70 kDa

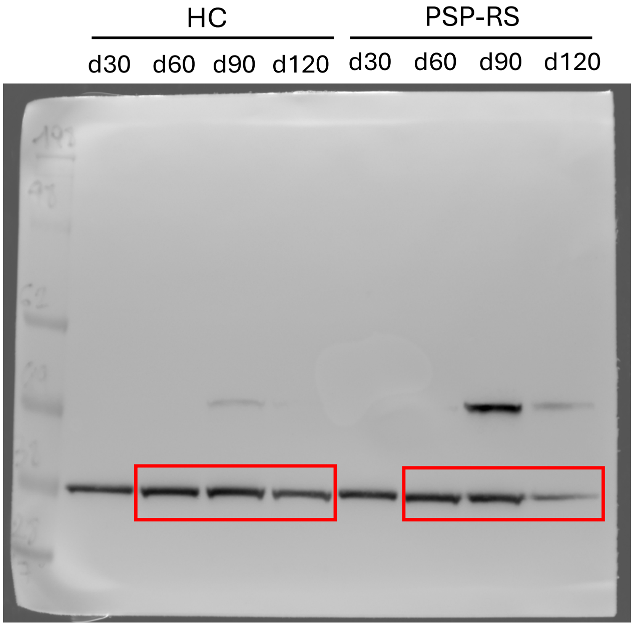


GAPDH MW 36 kDa

### 8) Figure 4b

1° replicate

d120

HC PSP-RS

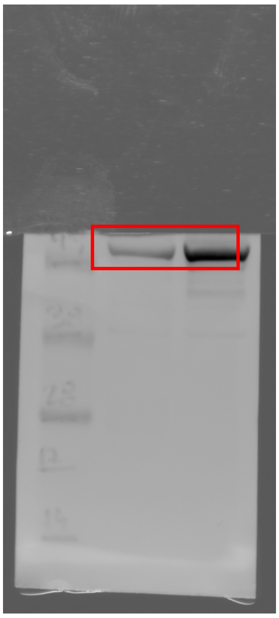

GFAP MW 50 kDa

d120

HC PSP-RS

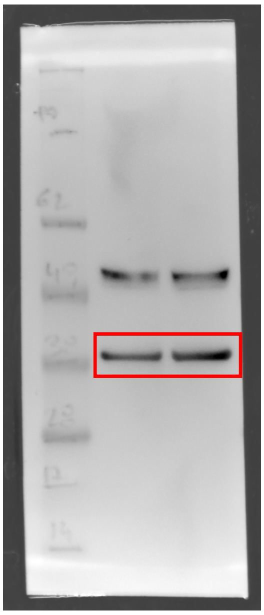

GAPDH MW 36 kDa

2° replicate

HC PSP-RS

d120 d120

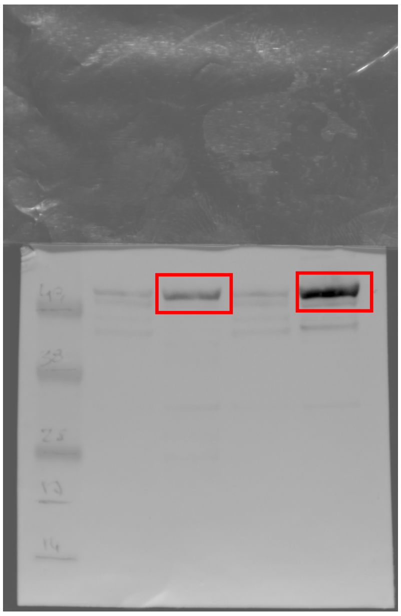

GFAP MW 50 kDa

HC PSP-RS

d120 d120

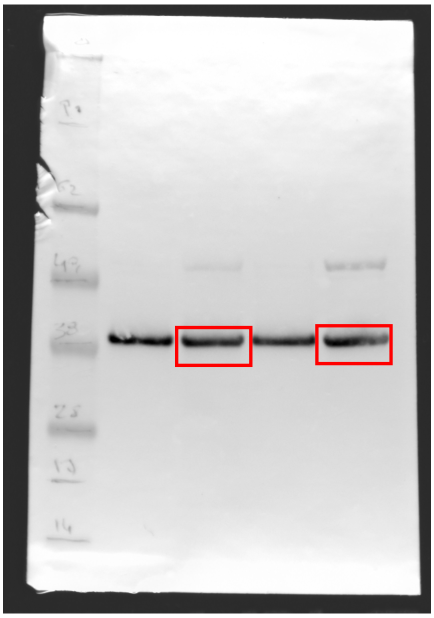

GAPDH MW 36 kDa
